# Serum preprocessing workflows differentially shape biological readout in data-independent acquisition proteomics of systemic juvenile idiopathic arthritis

**DOI:** 10.64898/2026.08.15.745022

**Authors:** Hironori Sato, Shinji Akioka, Ryo Konno, Yusei Okuda, Osamu Ohara, Yusuke Kawashima

## Abstract

Serum proteomics is increasingly used for minimally invasive biomarker discovery and disease phenotyping, and the choice of serum preprocessing workflow can shape proteome depth, quantitative characteristics, and downstream biological readouts. However, disease-oriented comparisons within a single cohort remain limited. Here, we compared four serum preprocessing workflows—Top14 depletion (TOP14D), tomato lectin affinity purification (TomAP), and two nanoparticle-based enrichment workflows (NPA and NPB)—using serum from six patients with systemic juvenile idiopathic arthritis (sJIA) and six age- and sex-matched healthy controls, and analyzed them using unified data-independent acquisition mass spectrometry (DIA-MS) and a statistical pipeline. We evaluated proteome depth, missingness, quantitative characteristics, group separation, differential abundance signatures, pathway enrichment, curated sJIA-related gene set coverage, pre-ranked gene set enrichment analysis (GSEA) results, and detection of inflammasome/interferon-related proteins. TomAP yielded the greatest proteome depth (7612 proteins), followed by NPB (6735 proteins) and NPA (6602 proteins), whereas TOP14D yielded the smallest protein set (3303 proteins). Principal component analysis (PCA) showed a separation between the sJIA and control groups for all workflows. Differentially expressed proteins (DEPs) showed limited overlap, with only 75 DEPs common to all four workflows. Functional enrichment patterns were workflow-dependent; TOP14D and TomAP mainly captured neutrophil/myeloid and inflammatory processes, whereas NPA and NPB captured RNA processing- and translation-related signals. TomAP showed relatively broad coverage and positive enrichment of curated sJIA-related gene sets associated with inflammation, innate immunity, and macrophage activation syndrome (MAS). Inflammasome/interferon-related proteins, including NLRC4, PYCARD, GSDMD, MEFV, IL-18, OAS3, and MYD88, showed workflow-dependent detectability and differential abundance. These findings support a disease-oriented benchmark for fit-for-purpose workflow selection according to the disease axis and analytical objective rather than proteome depth alone.

## Introduction

Human serum proteomics is widely used as a minimally invasive and readily accessible “liquid biopsy” technique to characterize disease phenotypes and identify candidate biomarkers (1, 2). In recent years, acquisition and analysis pipelines based on data-independent acquisition mass spectrometry (DIA-MS), which generally provides more reproducible quantitative measurements than data-dependent acquisition (DDA), have been increasingly adopted for large-scale cohort applications. DIA-MS improves data completeness, quantitative accuracy, and reproducibility through comprehensive acquisition (3). In parallel, advances in computational methods, including in silico spectral library generation and neural network-based analysis, have improved peptide and protein identification and quantification from DIA-MS data and facilitated the practical implementation of DIA-based workflows (4).

However, serum proteins span a wide and dynamic range, which remains a major analytical challenge. Approximately 20 highly abundant proteins, including albumin and immunoglobulins, limit the effective quantitative range of MS-based analyses, thereby hindering the detection and quantification of low-abundance proteins of physiological or pathological relevance (1, 5). To address this challenge, several serum preprocessing workflows have been proposed, including high-abundance protein depletion, nanoparticle-based enrichment designed to compress and redistribute the serum protein composition, and affinity enrichment strategies that emphasize specific subproteomes. Immunoaffinity depletion is a classical and readily implementable approach that can increase protein identification in the low-abundance range; however, it may also introduce co-depletion and workflow-specific quantitative bias (6). Nanoparticle-based enrichment using the protein corona can redistribute the circulating proteome and recover a detected protein set that differs from that obtained using conventional workflows (7). Lectin affinity enrichment emphasizes glycosylated secreted and membrane-associated proteins and can extend the classes of proteins detectable in serum along a distinct biological axis (8, 9). Additionally, antibody- and aptamer-based high-multiplex proteomic assays, such as Olink and SomaScan, have been rapidly adopted (10, 11). Thus, in serum proteomics, the choice of serum preprocessing workflow itself, in addition to the acquisition and analysis methods, can influence proteome depth, missingness, quantitative characteristics, emphasized protein classes, and biological readouts. Although comparative studies on depletion and enrichment workflows have increased in recent years, many have been technical benchmarks based on healthy donors or replicates, with a primary focus on proteome depth, repeatability, quantification bias, and contamination. Consequently, limited evidence is available from studies that have evaluated, within the same disease and cohort, how workflow-dependent differences are linked to the detection of specific disease axes using a unified DIA-MS pipeline (12). Recent workflow comparisons in plasma proteomics have emphasized that increased proteome depth does not necessarily translate into quantitative robustness or biological fidelity and that the effects of preanalytical variation and cellular carry-over require workflow-specific evaluation, particularly for nanoparticle-based enrichment (6). The discussions, however, have largely relied on plasma-based technical benchmarks. Consequently, how workflow-dependent differences relate to the detection of specific disease axis (when the same disease cohort is profiled in serum under a unified DIA-MS pipeline) remains insufficiently evaluated.

Systemic juvenile idiopathic arthritis (sJIA) is a childhood-onset autoinflammatory disease characterized by systemic inflammation, including remittent fever, rash, and arthritis (13, 14). The disease can become severe when complicated by macrophage activation syndrome (MAS), a secondary hemophagocytic syndrome. In sJIA/MAS, elevated IL-18 levels are associated with disease activity and MAS risk, and the potential role of IL-18 as an aid to diagnosis, stratification, and therapeutic targeting has been discussed (15, 16). In addition, an axis involving IFN-γ–related chemokines, such as CXCL9, has been reported to be associated with MAS, and the utility of CXCL9 as a circulating marker has been investigated (17). Subtypes based on inflammasome-related gene expression, such as *MYD88*-dominant and *NLRC5*-dominant groups, have also been reported (18). These findings indicate the importance of innate immunity, inflammasomes, and IFN-related signaling in sJIA, while also suggesting the need for stratification that accounts for molecular heterogeneity. Therefore, sJIA serum represents a suitable disease-oriented benchmark for evaluating the disease axes that are preferentially captured by different serum preprocessing workflows.

In this study, we compared four serum preprocessing workflows—Top14 depletion (TOP14D), tomato lectin affinity purification (TomAP), and two nanoparticle-based enrichment workflows (NPA and NPB)—using serum from the same sJIA cohort and a unified DIA-MS acquisition and statistical analysis pipeline. We comprehensively evaluated (i) proteome depth, missingness, quantitative range, reproducibility, sample correlation, and group separation; (ii) differential abundance analysis between sJIA and control samples, together with pathway-level characteristics; and (iii) the detectability of candidate proteins centered on inflammasome/interferon-related axes.

The aim of this study was not to rank serum preprocessing workflows on a single scale but to provide practical guidance for fit-for-purpose workflow selection according to the analytical objective.

### Experimental procedures

#### Study design and participants

We conducted a cross-sectional study of patients with sJIA treated at the Kyoto Prefectural University of Medicine Hospital. The study was conducted in accordance with the principles of the Declaration of Helsinki and was approved by the Ethics Review Board of Kyoto Prefectural University of Medicine (approval numbers: ERB-C-385-2, ERB-C-1613-2, and ERB-C-3977). Written informed consent was obtained from study participants and/or their legal guardians. Eligible patients were outpatients who had been diagnosed with sJIA according to the International League of Associations for Rheumatology (ILAR) classification criteria (13, 19). Serum samples were obtained at the initial visit within 2 weeks of disease onset or during active, early-stage disease (n = 6). Controls were age- and sex-matched children without inflammation attributable to other rheumatic diseases, acute infection, surgery, trauma, malignancy, or other causes (n = 6). The clinical characteristics of the study participants are summarized in Table 1.

**Table 1.**
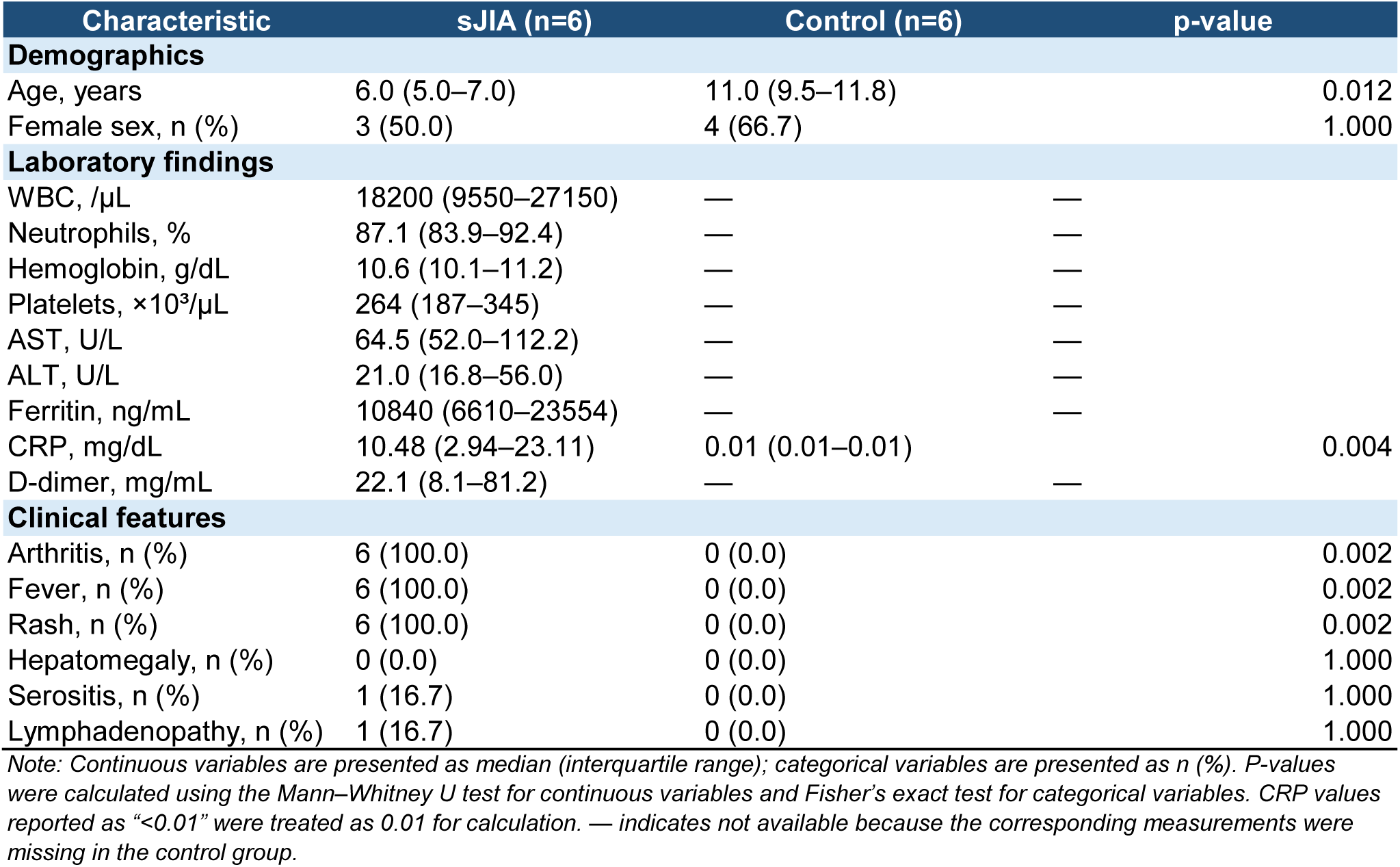
Clinical characteristics of patients with sJIA and healthy controls.

| Characteristic | sJIA (n=6) | Control (n=6) | p-value |
| --- | --- | --- | --- |
| **Demographics** |  |  |  |
| Age, years | 6.0 (5.0–7.0) | 11.0 (9.5–11.8) | 0.012 |
| Female sex, n (%) | 3 (50.0) | 4 (66.7) | 1.000 |
| **Laboratory findings** |  |  |  |
| WBC, /μL | 18200 (9550–27150) | — | — |
| Neutrophils, % | 87.1 (83.9–92.4) | — | — |
| Hemoglobin, g/dL | 10.6 (10.1–11.2) | — | — |
| Platelets, $\times 10^3/\mu$L | 264 (187–345) | — | — |
| AST, U/L | 64.5 (52.0–112.2) | — | — |
| ALT, U/L | 21.0 (16.8–56.0) | — | — |
| Ferritin, ng/mL | 10840 (6610–23554) | — | — |
| CRP, mg/dL | 10.48 (2.94–23.11) | 0.01 (0.01–0.01) | 0.004 |
| D-dimer, mg/mL | 22.1 (8.1–81.2) | — | — |
| **Clinical features** |  |  |  |
| Arthritis, n (%) | 6 (100.0) | 0 (0.0) | 0.002 |
| Fever, n (%) | 6 (100.0) | 0 (0.0) | 0.002 |
| Rash, n (%) | 6 (100.0) | 0 (0.0) | 0.002 |
| Hepatomegaly, n (%) | 0 (0.0) | 0 (0.0) | 1.000 |
| Serositis, n (%) | 1 (16.7) | 0 (0.0) | 1.000 |
| Lymphadenopathy, n (%) | 1 (16.7) | 0 (0.0) | 1.000 |

*Note:* Continuous variables are presented as median (interquartile range); categorical variables are presented as n (%). P-values were calculated using the Mann–Whitney U test for continuous variables and Fisher's exact test for categorical variables. CRP values reported as "<0.01" were treated as 0.01 for calculation. — indicates not available because the corresponding measurements were missing in the control group.

#### Serum proteomics

Serum samples were processed using three preprocessing strategies, which yielded four analytical workflows. First, serum was processed using the tomato lectin affinity purification method (TomAP) with the Maelstrom 9610 instrument, as previously described (9). Briefly, 25 µL of streptavidin bead suspension (Cytiva, Marlborough, MA, USA) was added to 600 µL of protein-free blocking buffer (Setsuyaku-Kun Supporter, DRC, Tokyo, Japan) diluted 10-fold with Tris-buffered saline (TBS). Next, 10 µL of 2 µg/µL tomato lectin (Vector Laboratories, Burlingame, CA, USA) was added. The mixture was gently agitated for 30 min, and the beads were washed once with 1.2 mL of dilution/wash buffer consisting of TBS containing 0.0005% Tween 20. Serum diluted in dilution/wash buffer was then added to the beads by mixing 100 µL of serum with 400 µL of dilution/wash buffer, followed by incubation for 60 min with gentle mixing. The beads were washed twice with 1.2 mL of dilution/wash buffer and rinsed with 1.2 mL TBS. For protein elution, the beads were mixed with 70 µL of 0.5% TFA containing 0.05% LMNG for 15 min. After magnetic separation, the eluate was collected and neutralized by adding 500 mM Tris-HCl (pH 8.0) containing 20 mM CaCl to adjust the pH to approximately 8.0. For digestion, 500 ng of Trypsin/Lys-C Mix (cat. no. V5072; Promega) was added to the neutralized eluate and the mixture was incubated at 37 °C for 16 h. The resulting peptides were reduced and alkylated by adding 14 µL of a solution containing 110 mM tris(2-carboxyethyl)phosphine and 440 mM 2-chloroacetamide, followed by incubation at 80 °C for 15 min. The samples were then acidified with 35 µL 5% TFA. The peptides were desalted using GL-Tip SDB tips (GL Sciences, Tokyo, Japan) according to the manufacturer’s instructions, eluted with 34% acetonitrile in 0.1% TFA, and dried using a centrifugal evaporator. The dried peptides were reconstituted in 0.02% DMNG containing 0.1% TFA.

Next, antibody column depletion was performed using Top14 Abundant Protein Depletion Mini Spin Columns (Thermo Fisher Scientific, Waltham, MA, USA) according to the manufacturer’s instructions (TOP14D method). Serum samples treated with TOP14D were subjected to clean-up and digestion using the SP3-LASP method with the Maelstrom 9610 instrument (20). Third, the serum samples were pretreated and digested into peptides using the Seer Proteograph XT, as previously described (6). Each sample was incubated with two nanoparticle mixtures to generate two fractions (NPA and NPB) per sample.

The digested peptides were injected directly onto a 75 µm × 30 cm nanoLC column (ReproSil-Pur C18, particle size 1.5 µm, 100 Å; CoAnn Technologies, Richland, WA, USA) at 60 °C, and separated using an 84.5-min gradient (A = 0.1% formic acid in water, B = 0.1% formic acid in 80% acetonitrile) consisting of 1% B with a flow rate of 600 μL/min in 0–0.5 min, 1–6% B with a flow rate of 600–200 μL/min in 0.5–4.5 min, 6–24% B with a flow rate of 200 μL/min in 4.5–60 min, 24–40% B with a flow rate of 200 μL/min in 60–78.5 min, 40–98% B with a flow rate of 200 μL/min in 78.5–79.5 min, 98% B with a flow rate of 200 μL/min in 79.5–80.5 min, 98% B with a flow rate of 200–600 μL/min in 80.5–81.5 min, 98% B with a flow rate of 600–750 μL/min in 81.5–82.5 min, and 98% B with a flow rate of 750 μL/min in 82.5–84.5 min. The peptides eluted from the column were analyzed using an Orbitrap Astral equipped with an InSpIon system (21). MS1 spectra were collected over m/z 380–980 at a resolution of 240,000 using the Orbitrap, with an automatic gain control target of 300% and a maximum injection time of 5 ms. MS2 spectra were collected at m/z 200–2,000 using the Astral analyzer, with an automatic gain control target of 300%, a maximum injection time of 3.5 ms, and a normalized collision energy of 25%. The isolation width for MS2 was set to 2 Th, with window placement optimization enabled.

The MS data were queried against an in silico human spectral library using DIA-NN v.1.9.2 (22). Initially, a spectral library was generated from the human protein sequence UniProt database (proteome ID UP000005640; 20,656 entries downloaded on November 1, 2024). The parameters for generating the spectral library were as follows: digestion enzyme, trypsin; missed cleavages, 2; peptide length range, 7–35; precursor charge range, 2–4; precursor mass range, 350–1,000; and fragment ion m/z range, 100–2,000. Additionally, “FASTA digest for library-free search/library generation,” “deep learning-based spectra, RTs and IM prediction,” “n-term M excision” were enabled. For the DIA-NN search, the following parameters were applied: MS1 accuracy of 10 ppm, MS2 accuracy of 10 ppm, protein inference based on genes, utilization of neural network classifiers in single-pass mode, quantification strategy using QuantUMS (high precision), cross-run normalization set to “RT-dependent.” Furthermore, “Unrelated runs,” “Peptideforms,” “Heuristic protein inference,” and “No shared spectra” were enabled. The threshold for protein identification was set at 1% or less for both the precursor and protein false discovery rates.

#### Experimental design and statistical rationale

The primary comparison was a two-group comparison between the sJIA and control groups within each serum preprocessing workflow. Comparisons between the workflows were performed as a disease-oriented benchmark within a single cohort. Because the workflows differed in their preprocessing design, including input amount, chemistry, and post-enrichment processing, the study was positioned not as a simple ranking of workflow performance but as a fit-for-purpose comparison according to the analytical objective.

For each workflow, we first evaluated the number of identified proteins, missingness, protein intensity distribution, coefficient of variation (CV), inter-sample correlation, and group separation using principal component analysis (PCA). We then independently performed differential abundance analysis between the sJIA and control groups for each workflow. For functional analysis, over-representation analysis was performed using proteins classified as upregulated in the sJIA group, and pre-ranked gene set enrichment analysis (GSEA) was conducted against curated sJIA-related gene sets. These analyses were used to compare workflow-dependent differences in the biological processes and pathways detected across the workflows.

#### Bioinformatics and statistical analysis

All downstream analyses were performed using RStudio 2025.09.2 (Posit). Data processing and visualization were performed primarily using tidyverse packages including dplyr, tidyr, readr, tibble, and ggplot2. Protein-level intensity matrices exported from DIA-NN were used for analysis, and intensity values of 0 or lower were considered missing. The resulting intensity values were log_2_-transformed before analysis.

For PCA, only proteins with finite values in at least 50% of the samples within each workflow were included, and the remaining missing values were imputed using protein-wise means. This imputation was limited to PCA and visualization purposes and was not used for differential abundance analysis. Two-group comparisons between the sJIA and control groups were performed using linear model estimation with the limma package and empirical Bayes moderation, together with peptide count-dependent variance adjustment using the DEqMS package. For each workflow, only proteins with log_2_ intensity values in at least three samples from both the sJIA and control groups were included in the differential abundance analysis. Model fitting was performed using lmFit/eBayes in limma, and SpectraCounteBayes in DEqMS was applied to the resulting statistics (23, 24). The Benjamini–Hochberg method was used for multiple testing corrections. Differentially expressed proteins (DEPs) were defined as proteins satisfying |log_2_FC| ≥ 0.58 (1.5-fold) and FDR < 0.05. Gene Ontology (GO) Biological Process enrichment analysis was performed using the enrichGO function in the ClusterProfiler package, and Reactome pathway analysis was performed using the enrichPathway function in the ReactomePA package. For both GO and Reactome analyses, p-values were adjusted for multiple testing using the Benjamini–Hochberg method, and the top pathways were plotted.

#### Construction of curated sJIA-related gene sets

To evaluate the key biological processes related to the pathophysiology of sJIA, we constructed curated sJIA-related gene sets comprising five categories: acute phase/complement, neutrophil activation, innate TLR/NLR signaling, cytokine signaling, and macrophage/MAS. Each category was generated by selecting terms and pathways related to the inflammatory pathophysiology of sJIA from the GO, Kyoto Encyclopedia of Genes and Genomes (KEGG), and Reactome databases. The terms, pathway IDs, and genes included in each gene set are listed in Tables S1–S5.

Data retrieval and ID conversion were performed using org.Hs.eg.db, GO.db, KEGGREST, ReactomePA, and reactome.db. Entries without UniProt IDs were excluded from the analysis, and the remaining entries were organized as curated gene sets, together with their corresponding gene symbols and source IDs.

GSEA of the curated gene sets was performed using the fgsea package. For each workflow, the log_2_FC values for each gene symbol obtained from the differential abundance analysis were used as the ranking score. For each gene set, fgsea was used to calculate the normalized enrichment score (NES) and FDR-adjusted p-value using the Benjamini–Hochberg method, and these values were compared across workflows.

## Results

### Comparison of identified proteins and quantitative performance across serum preprocessing workflows

First, we compared the number of proteins identified using each serum preprocessing workflow. TomAP identified the largest number of proteins (7612), whereas NPA (6602) and NPB (6735) also achieved high proteome depths. In contrast, TOP14D yielded the smallest protein set (3303 proteins; Fig. 1A). Comparison of the overlap among workflows revealed a group of proteins detected across all four workflows, as well as TomAP-specific proteins and proteins shared by NPA and NPB (Fig. 1B). These observations indicate that differences in serum preprocessing workflows are associated not only with the number of identified proteins but also with the composition of the detected protein set.

**Fig. 1.**
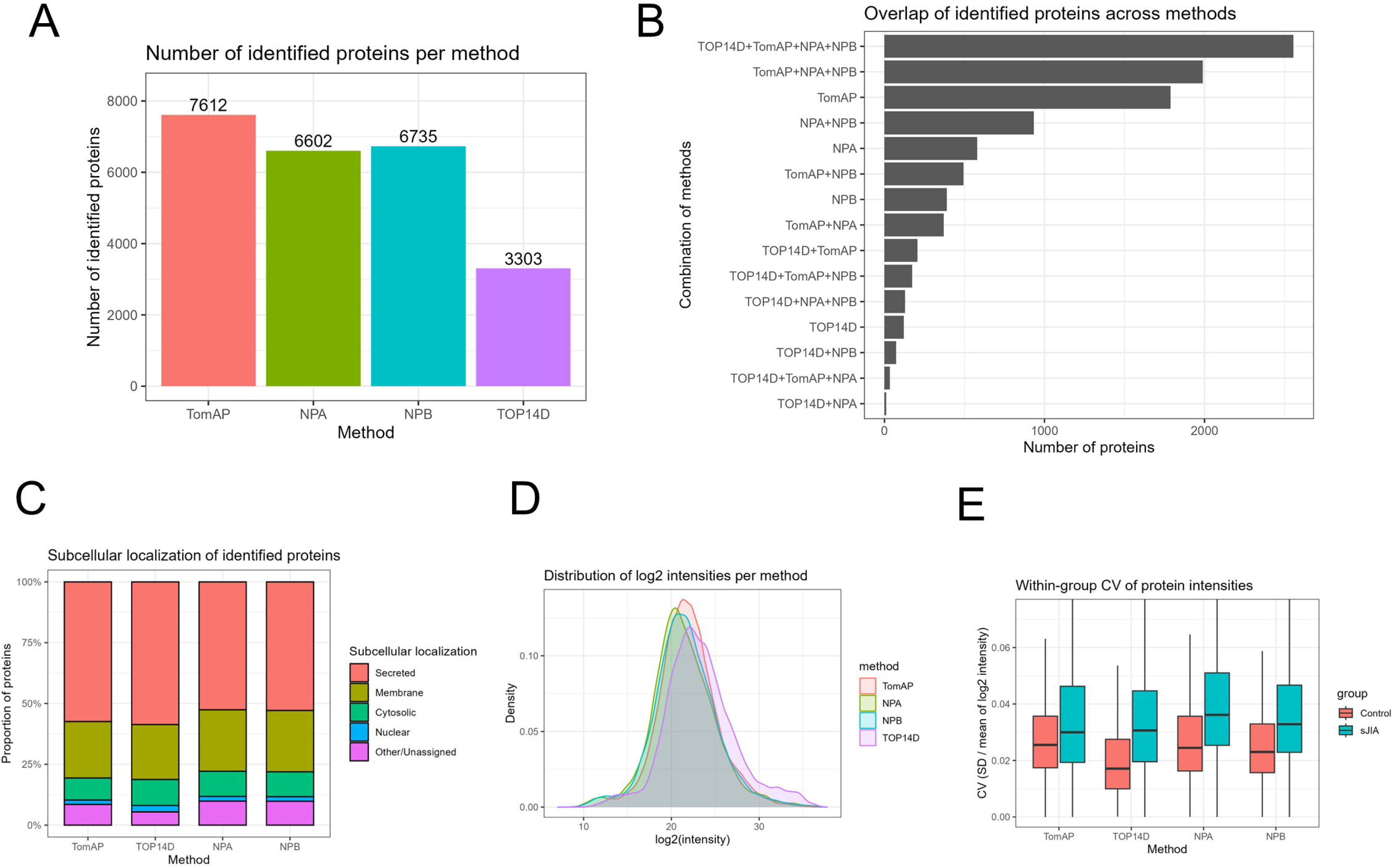
Comparison of proteome depth and quantitative characteristics across serum preprocessing workflows. A. Number of proteins identified by each serum preprocessing workflow: TomAP, NPA, NPB, and TOP14D. Values above the bars indicate the number of proteins identified in each workflow. B. Overlap of proteins identified across individual workflows and workflow combinations. Horizontal bars indicate the number of proteins corresponding to each combination of workflows shown on the left. C. Relative proportions of identified proteins classified according to subcellular localization. Proteins were categorized as Secreted, Membrane, Cytosolic, Nuclear, or Other/Unassigned. D. Density distributions of log_2_-transformed protein intensities for each workflow. E. Distributions of within-group coefficients of variation (CVs) of protein intensities in the control and sJIA groups for each workflow. CVs were calculated as the standard deviation divided by the mean of the log_2_-transformed protein intensities within each group. CV, coefficient of variation; NPA and NPB, nanoparticle-based enrichment workflows; sJIA, systemic juvenile idiopathic arthritis; TomAP, tomato lectin affinity purification; TOP14D, Top14 abundant protein depletion.

Next, we classified the identified proteins according to their subcellular localization and compared their proportions across the workflows. In all workflows, secreted proteins accounted for the largest proportion, consistent with the expected composition of serum-derived proteins (TomAP, 57%; TOP14D, 59%; NPA, 53%; and NPB, 53%), followed by membrane proteins, which accounted for approximately 25% (Fig. 1C). Cytosolic and nuclear proteins accounted for approximately 9–11% and 2–3%, respectively, and the proportions of these major categories were broadly similar across the workflows. In contrast, the proportion of other/unassigned proteins was higher in TomAP and NPA/NPB than in TOP14D, indicating workflow-dependent differences in the composition of these categories.

We then compared the quantitative characteristics of the protein-level data. Protein intensity distributions were broadly similar across the workflows, although differences were observed in the centers and spread of the distributions (Fig. 1D). Within-group CV distributions and inter-sample correlations showed broadly similar patterns across the workflows (Fig. 1E and Fig. S1A–D). PCA showed separation between the sJIA and control groups in all workflows (Fig. S1E–H). These descriptive analyses supported the use of each workflow for exploratory within-workflow comparisons but did not constitute a direct assessment of technical reproducibility.

### Differential protein signatures in sJIA differ across serum preprocessing workflows

Next, we examined how the detection of changes in sJIA-associated protein abundance differed across the serum preprocessing workflows. For each workflow, differential abundance analysis between the sJIA and control groups was performed independently, and the proteins included in the analysis and the differentially expressed proteins (DEPs) were identified (Table 2).

**Table 2.**
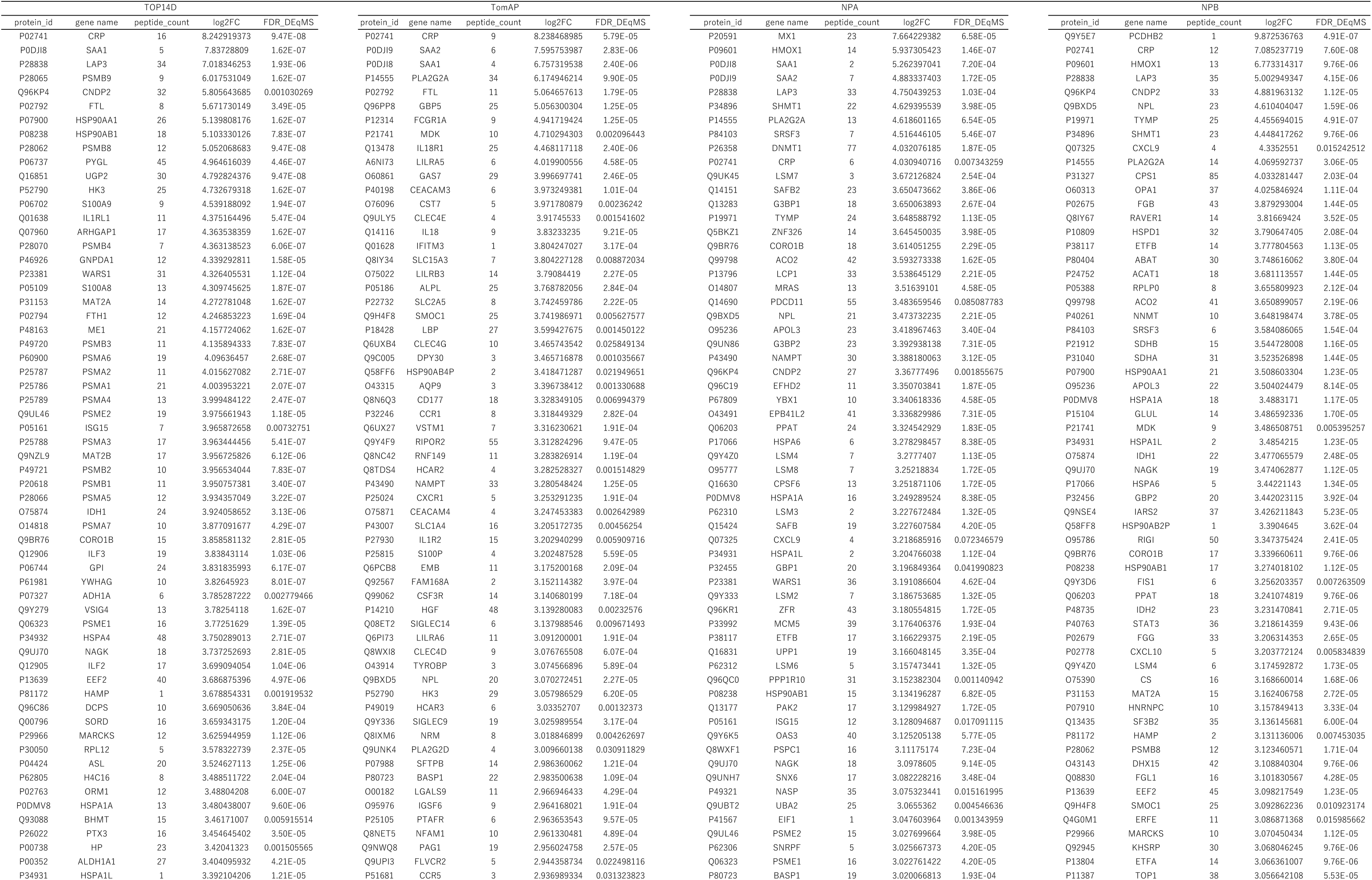
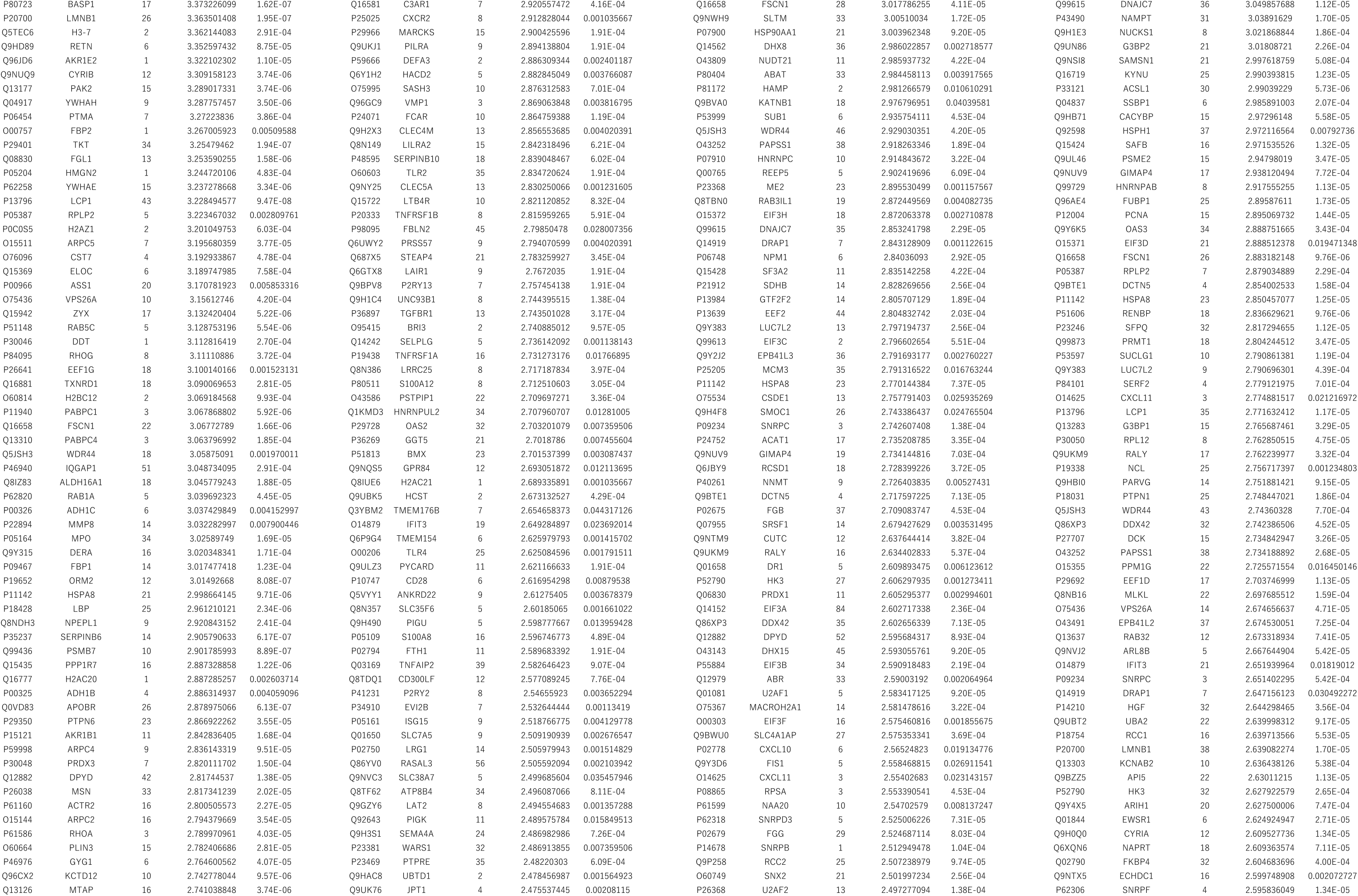
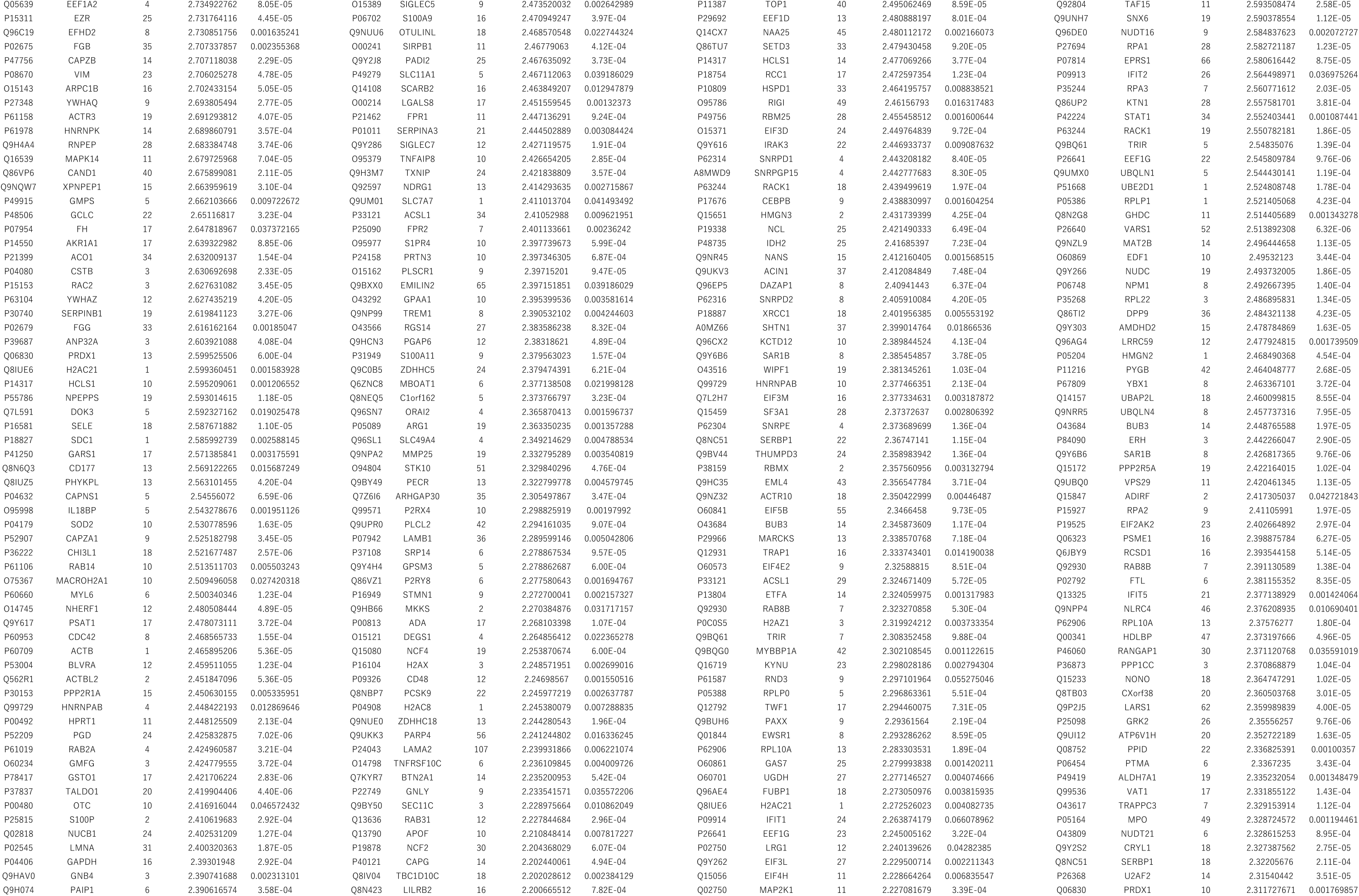
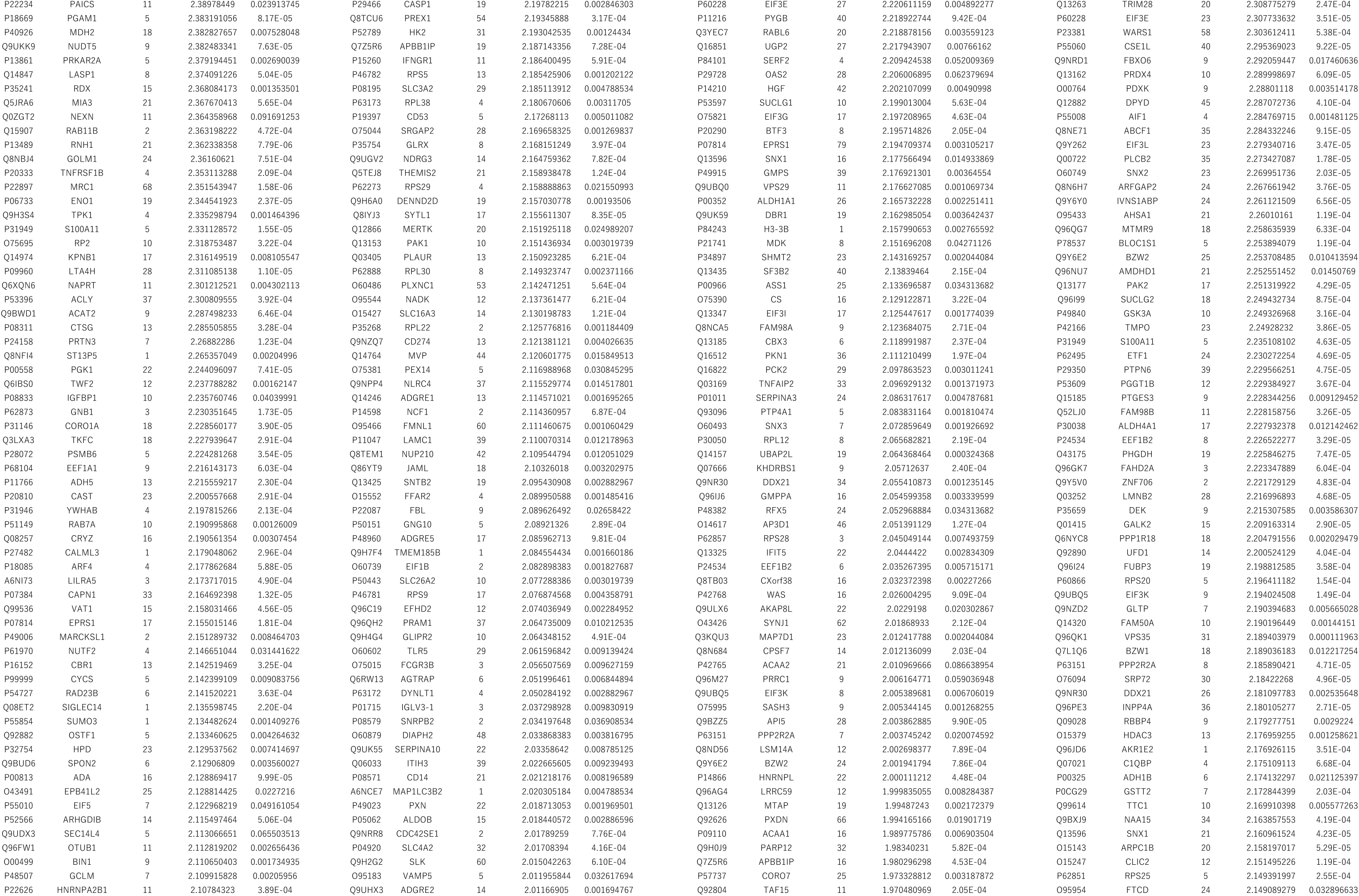
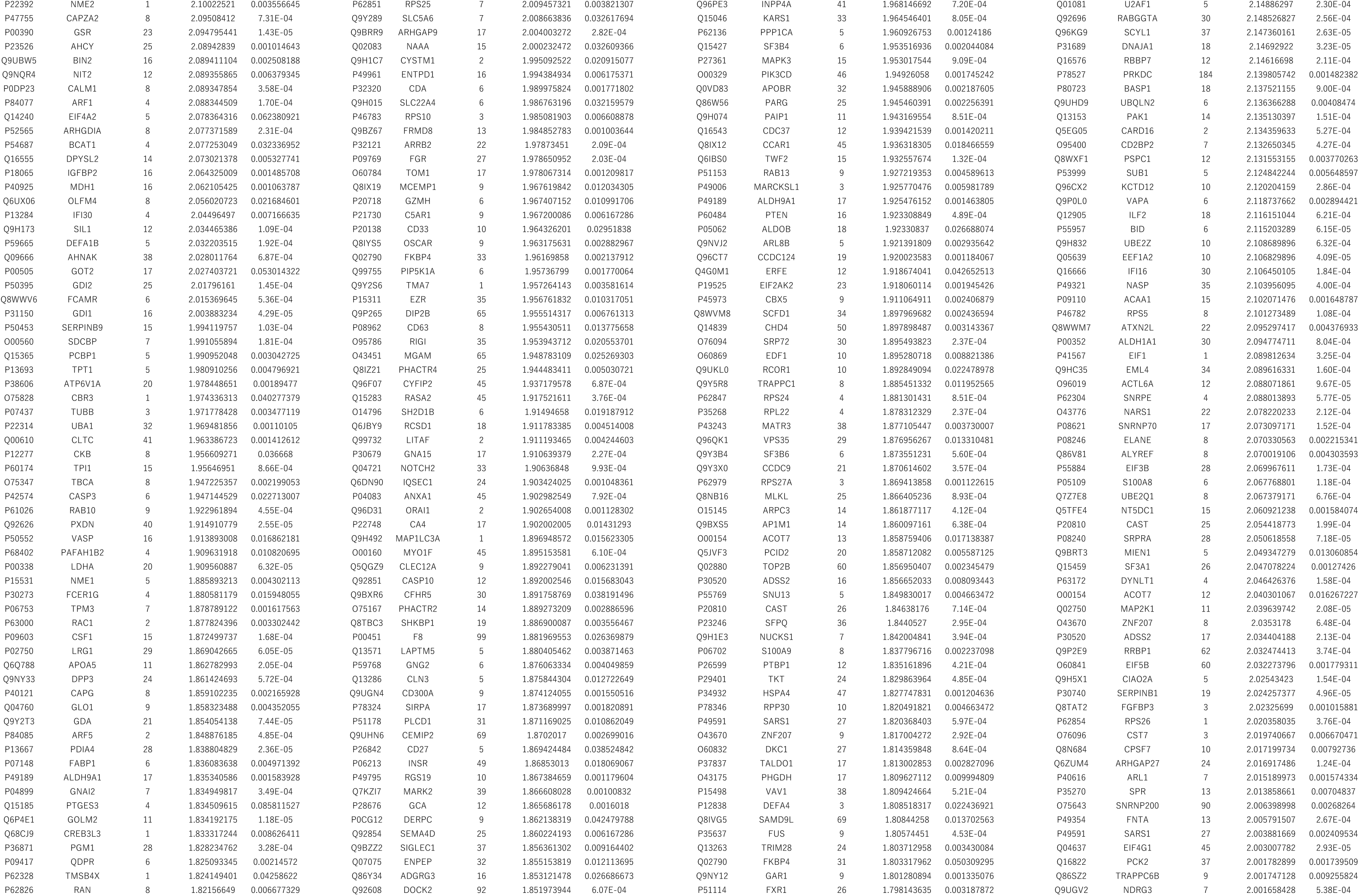
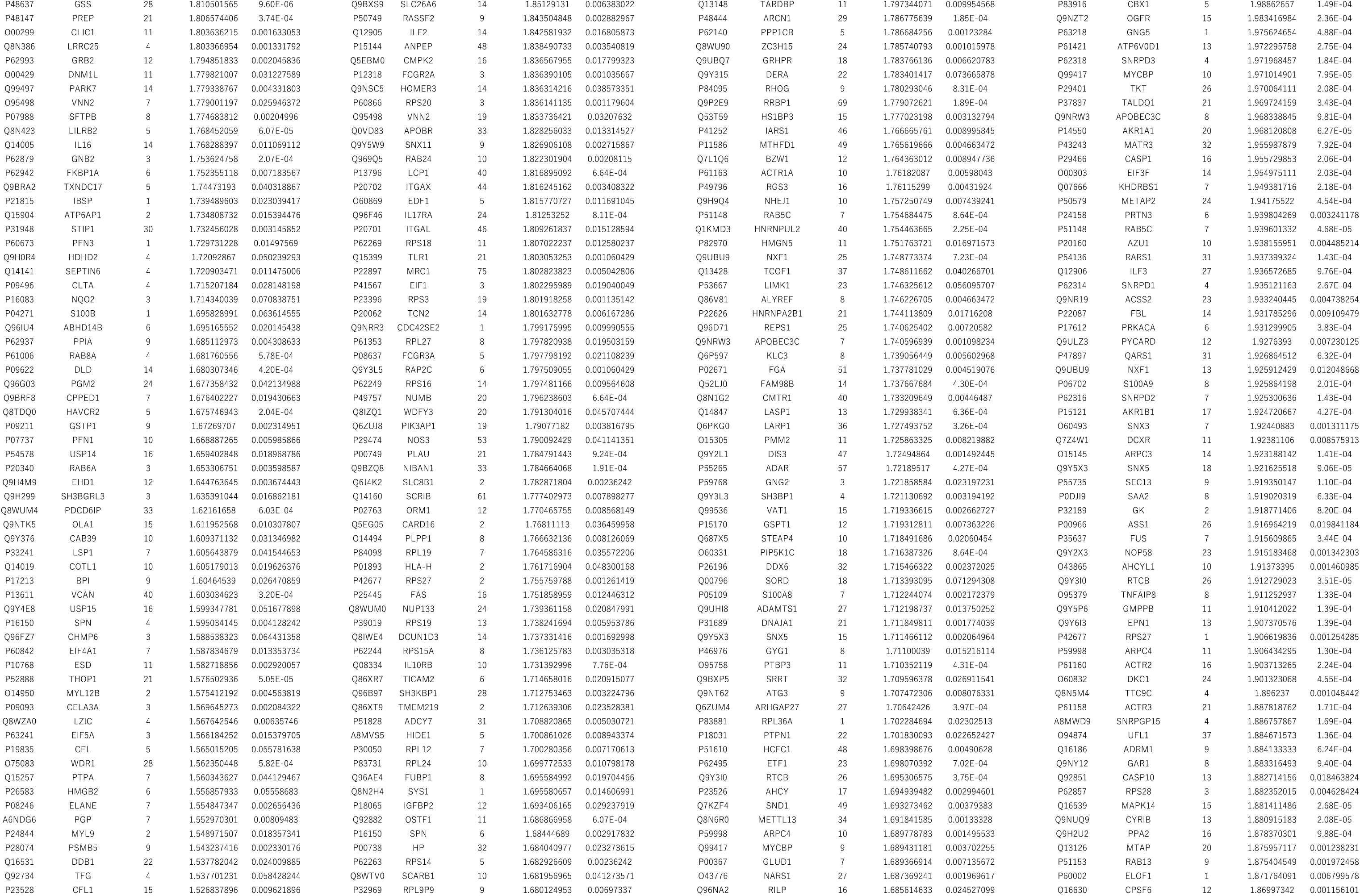
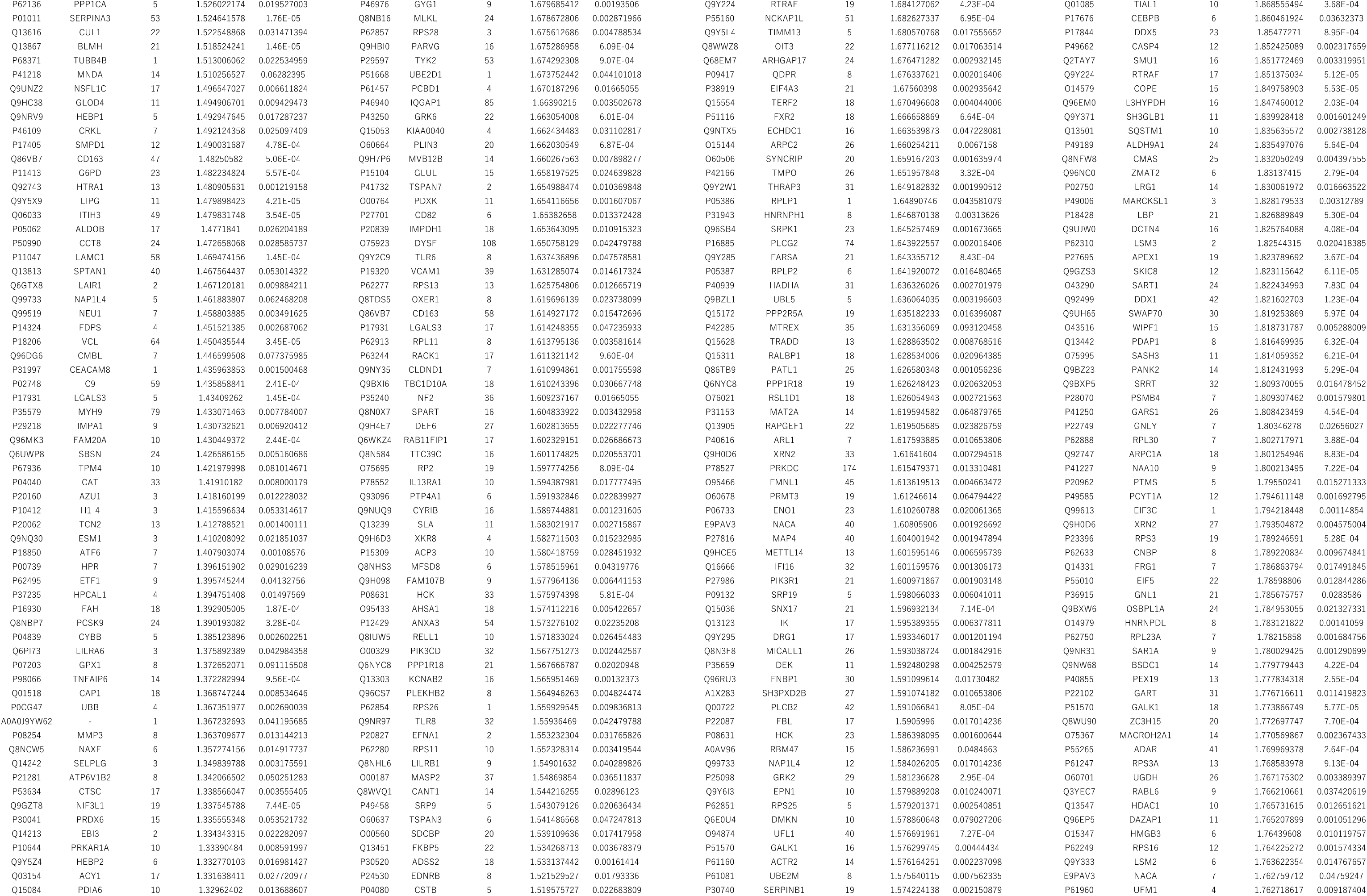
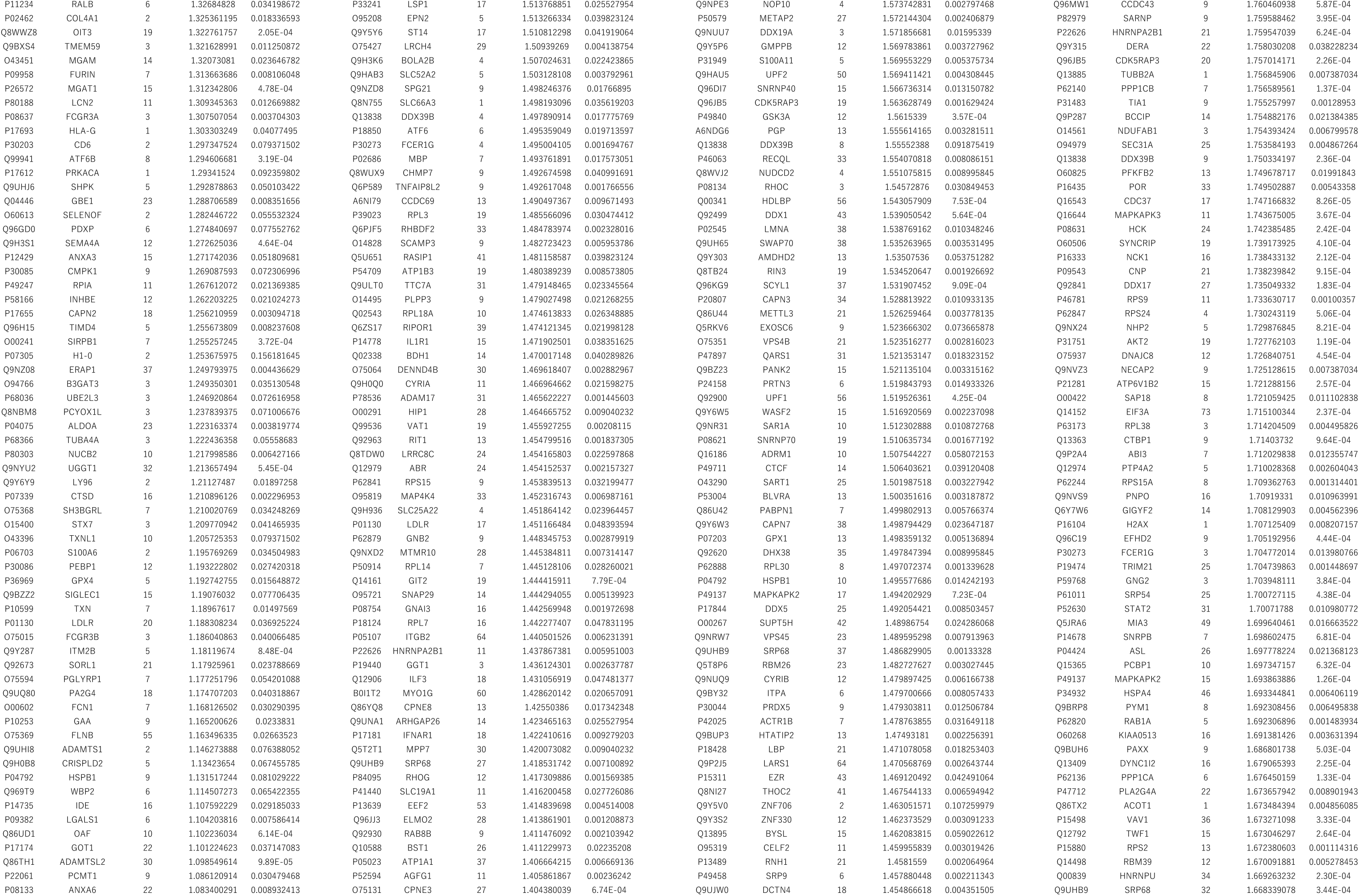
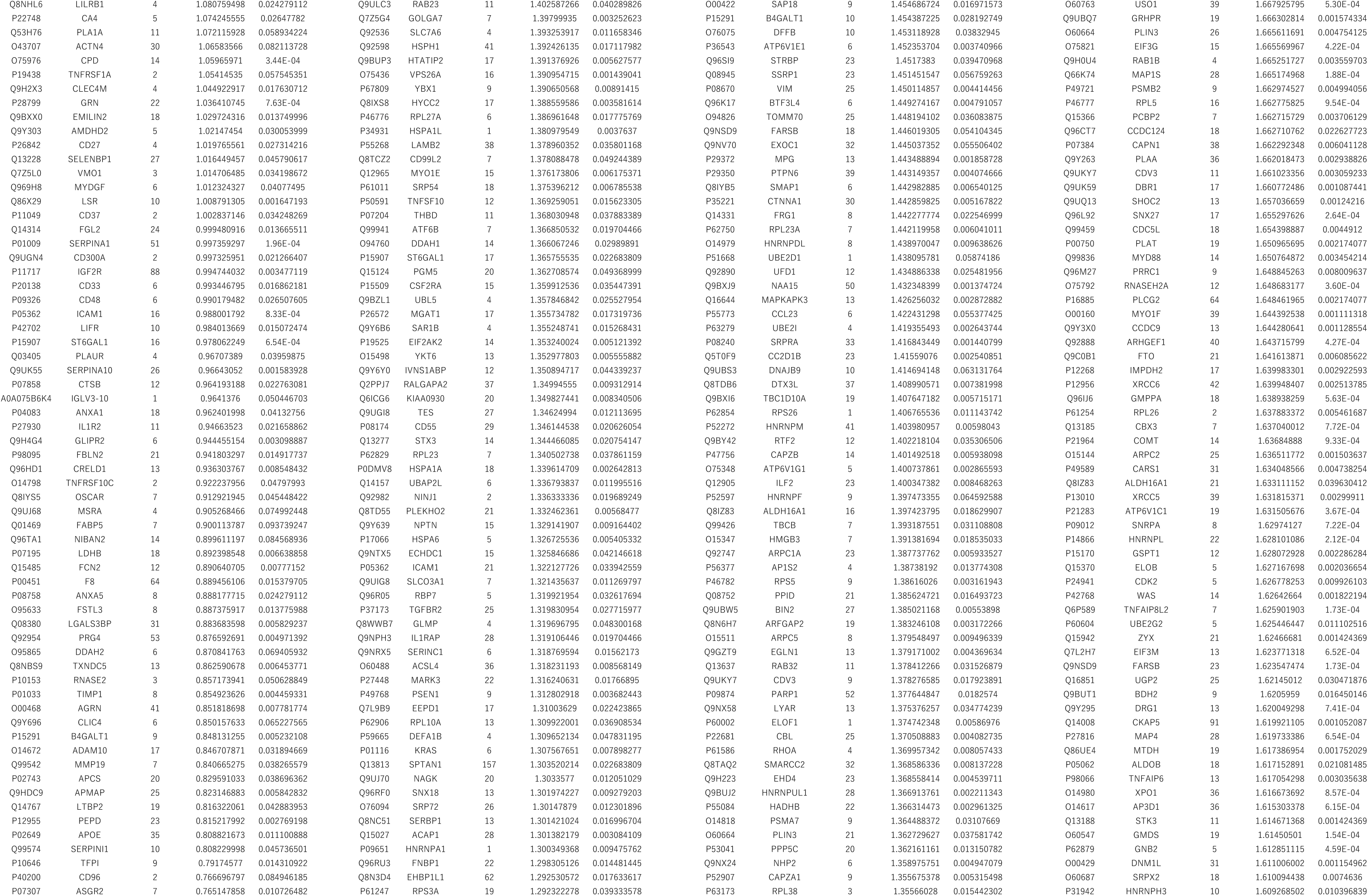
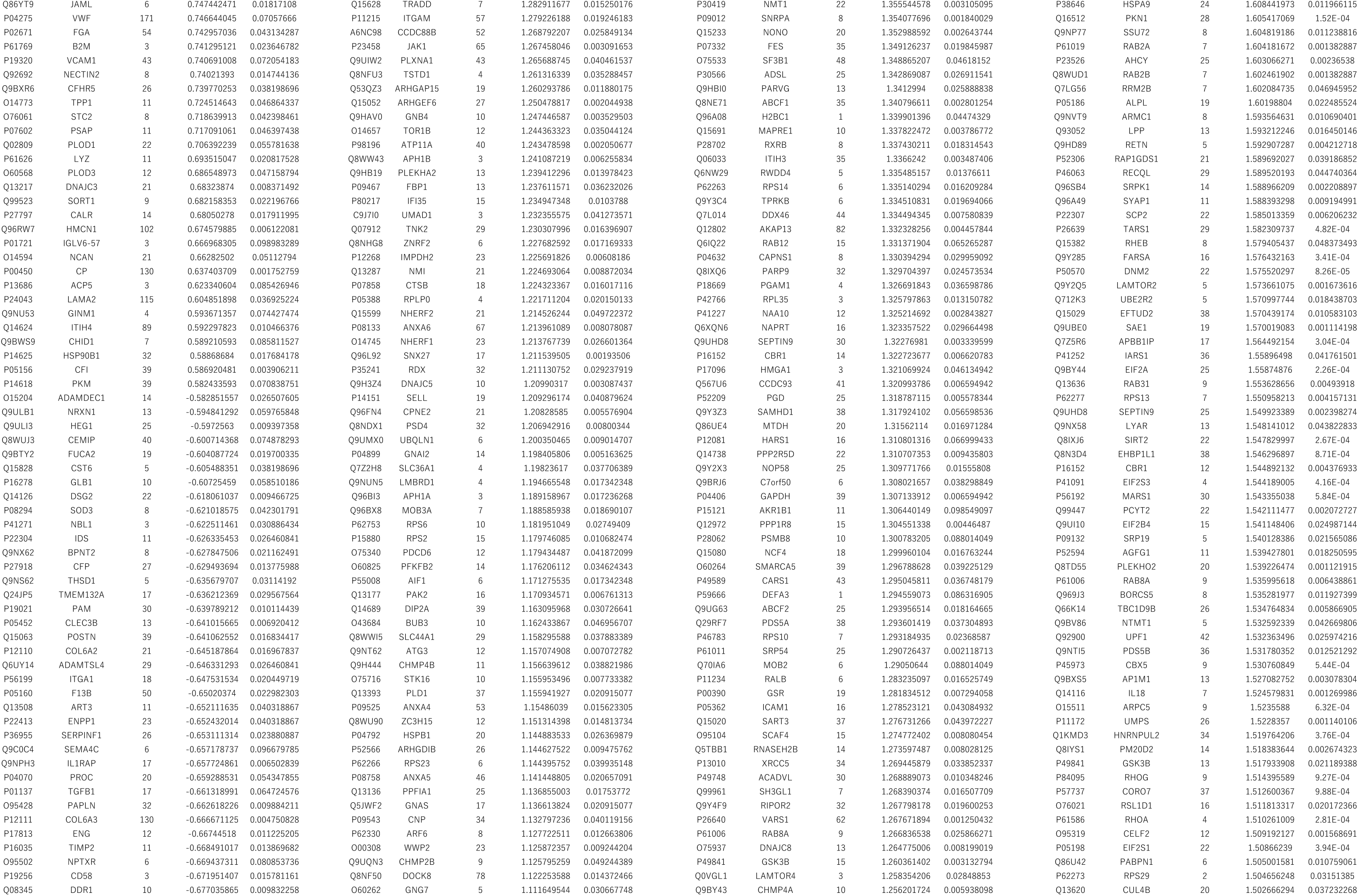
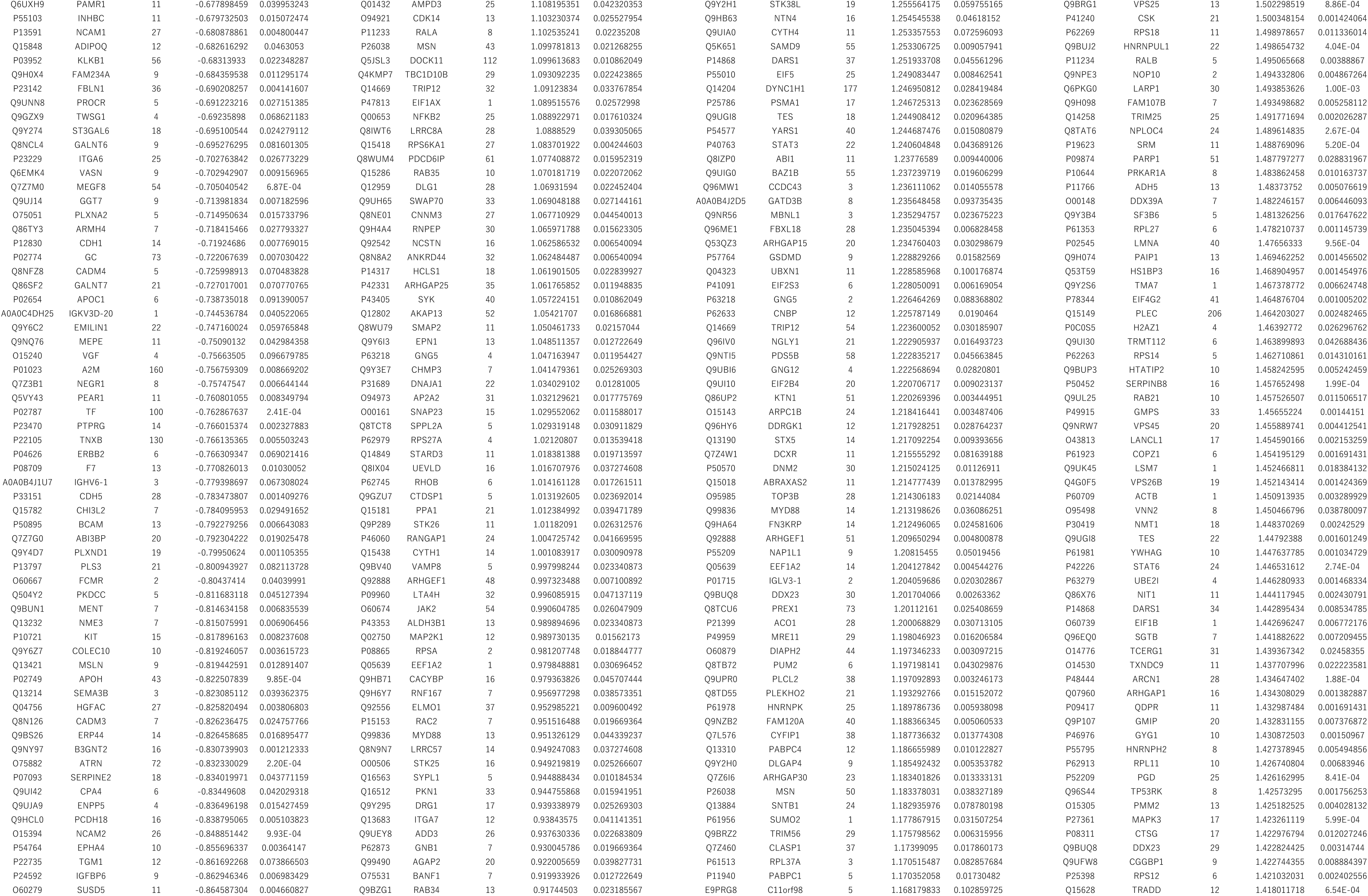
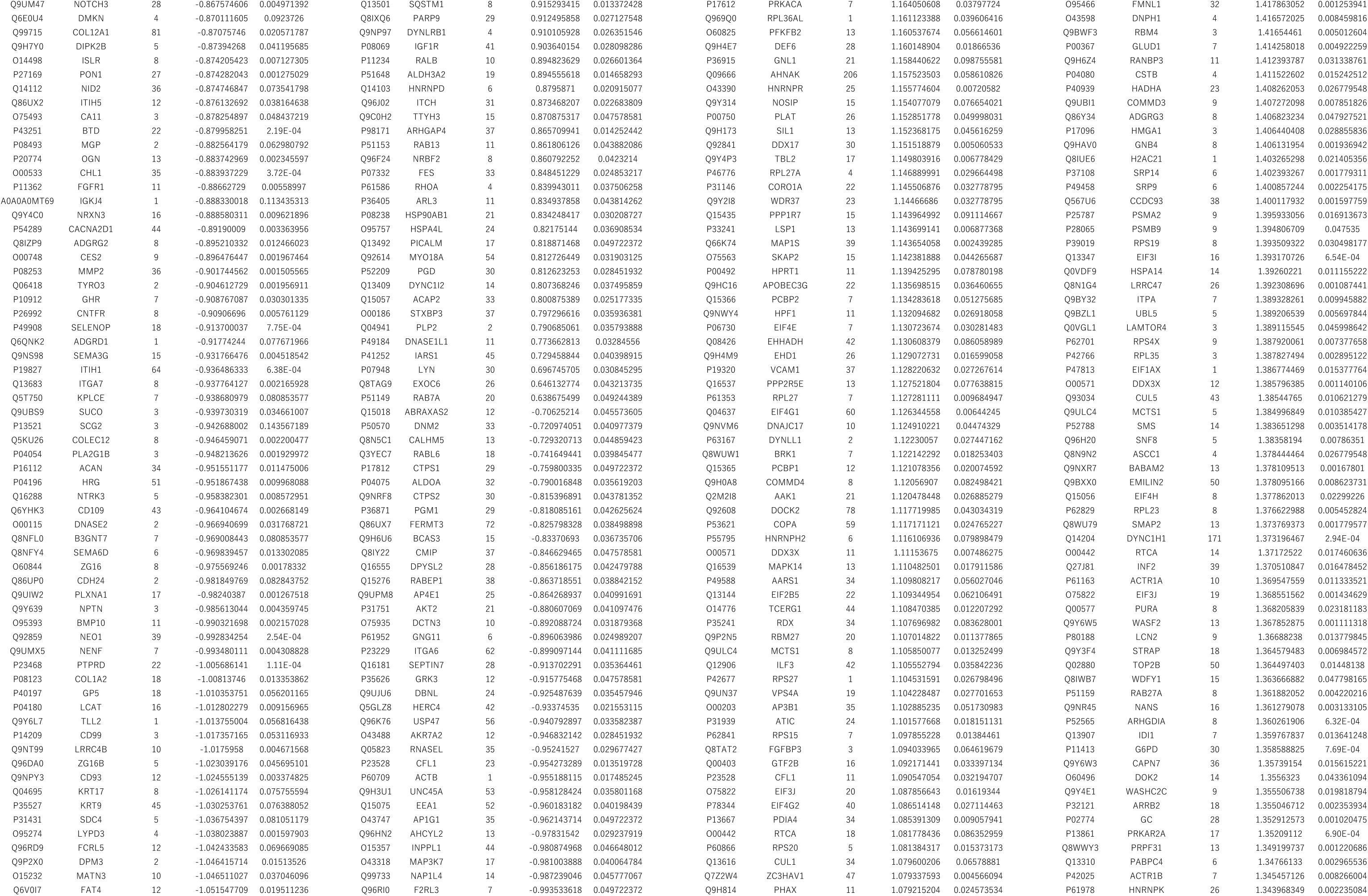
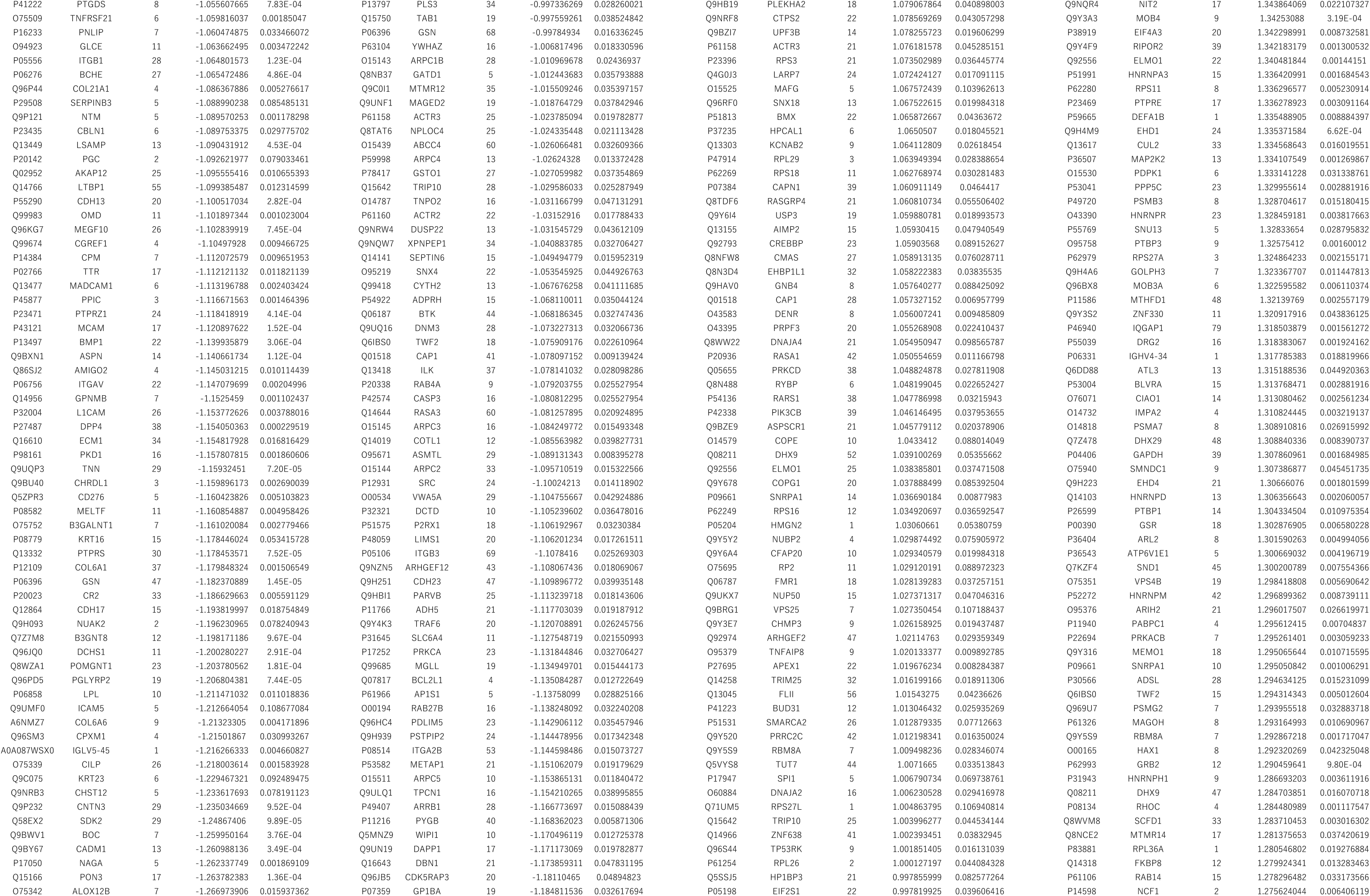
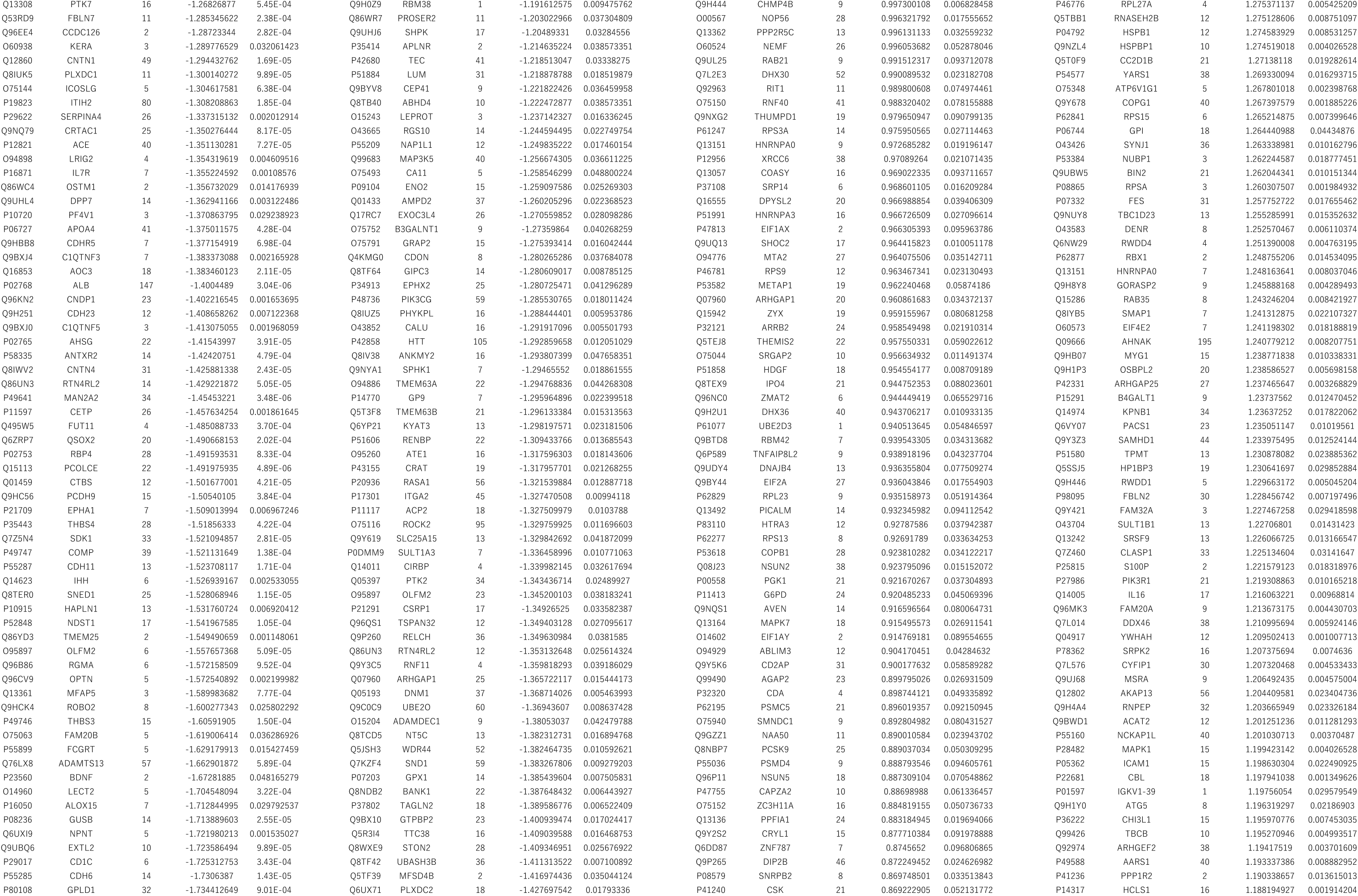
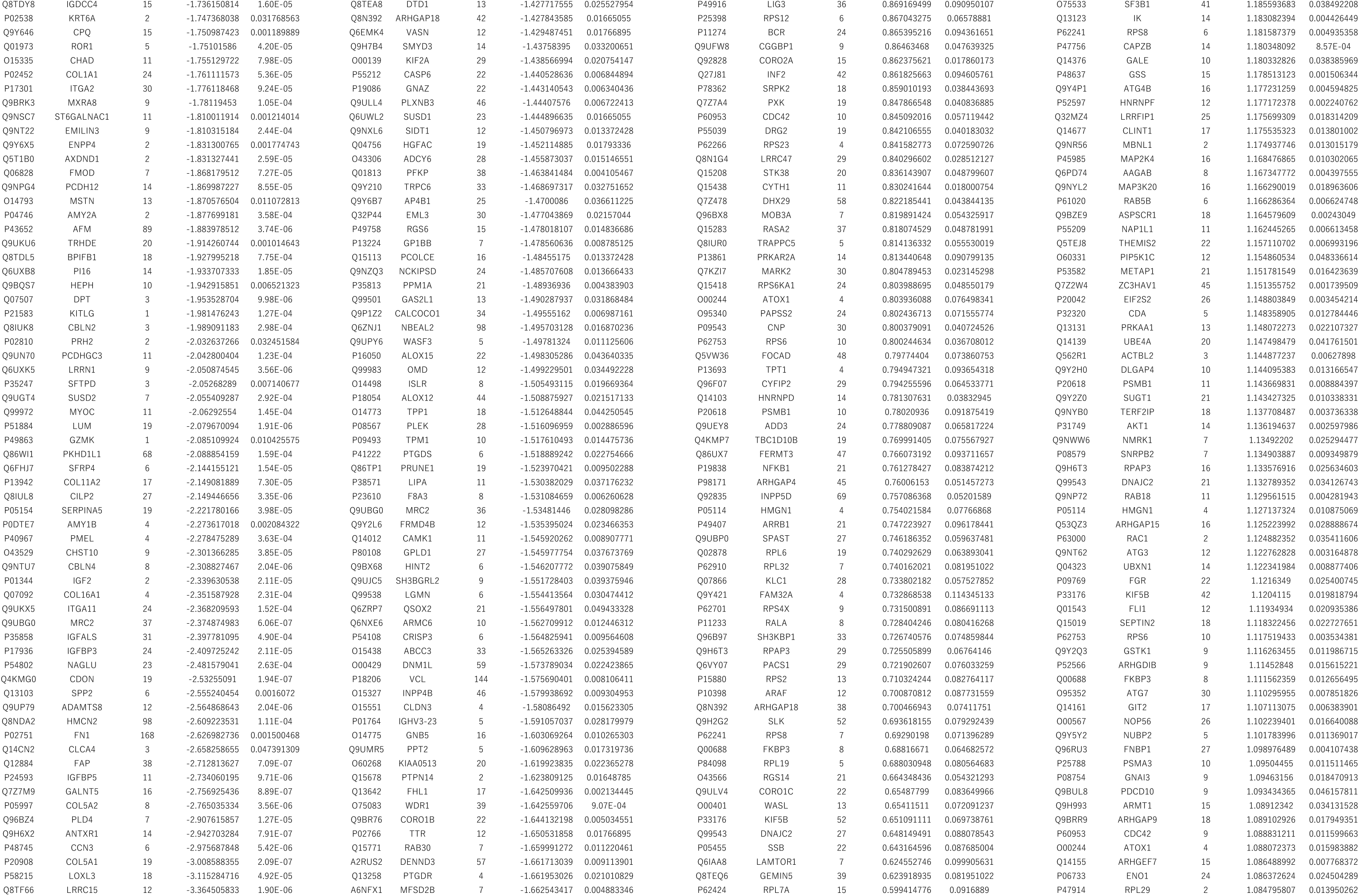
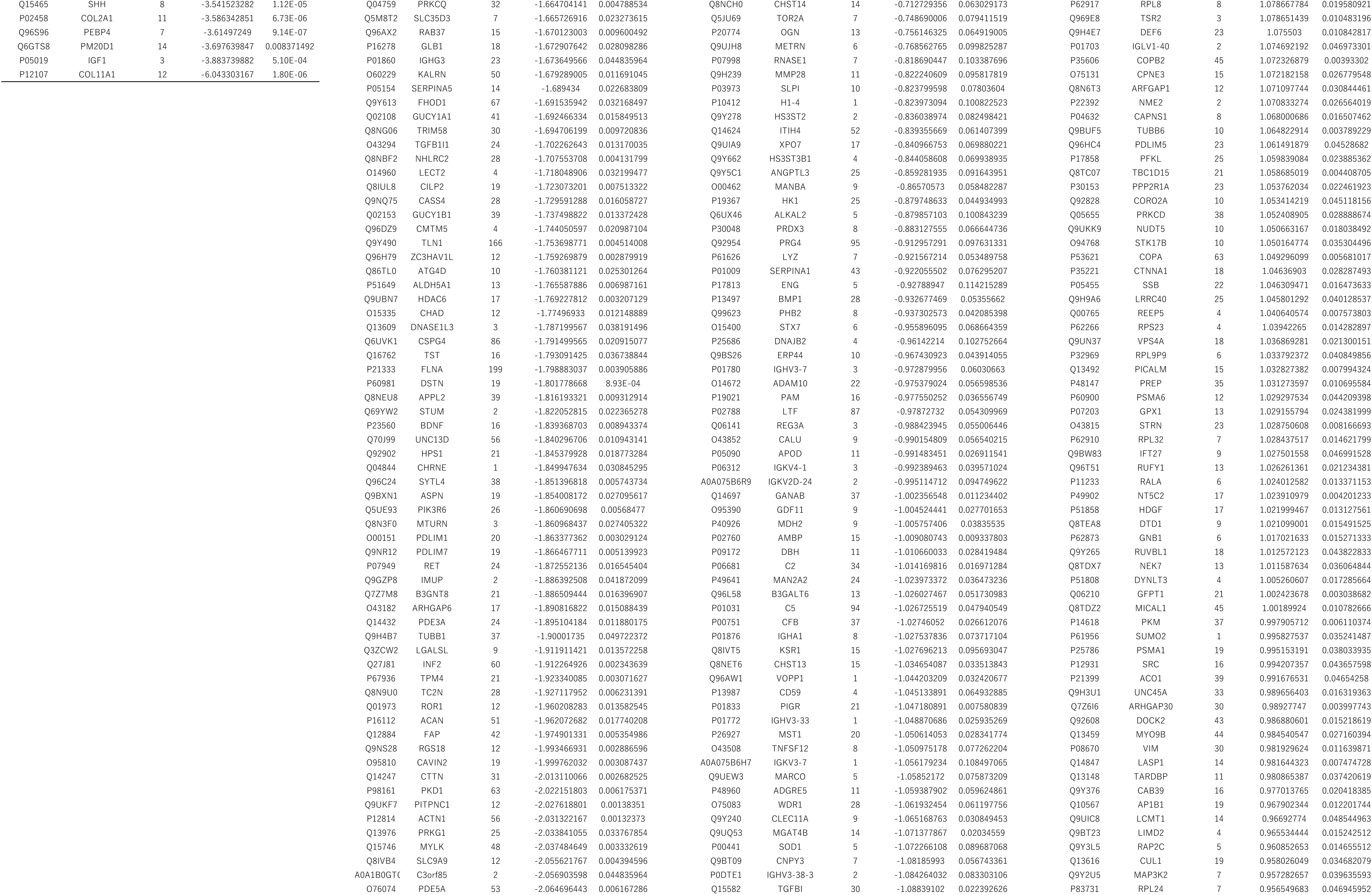
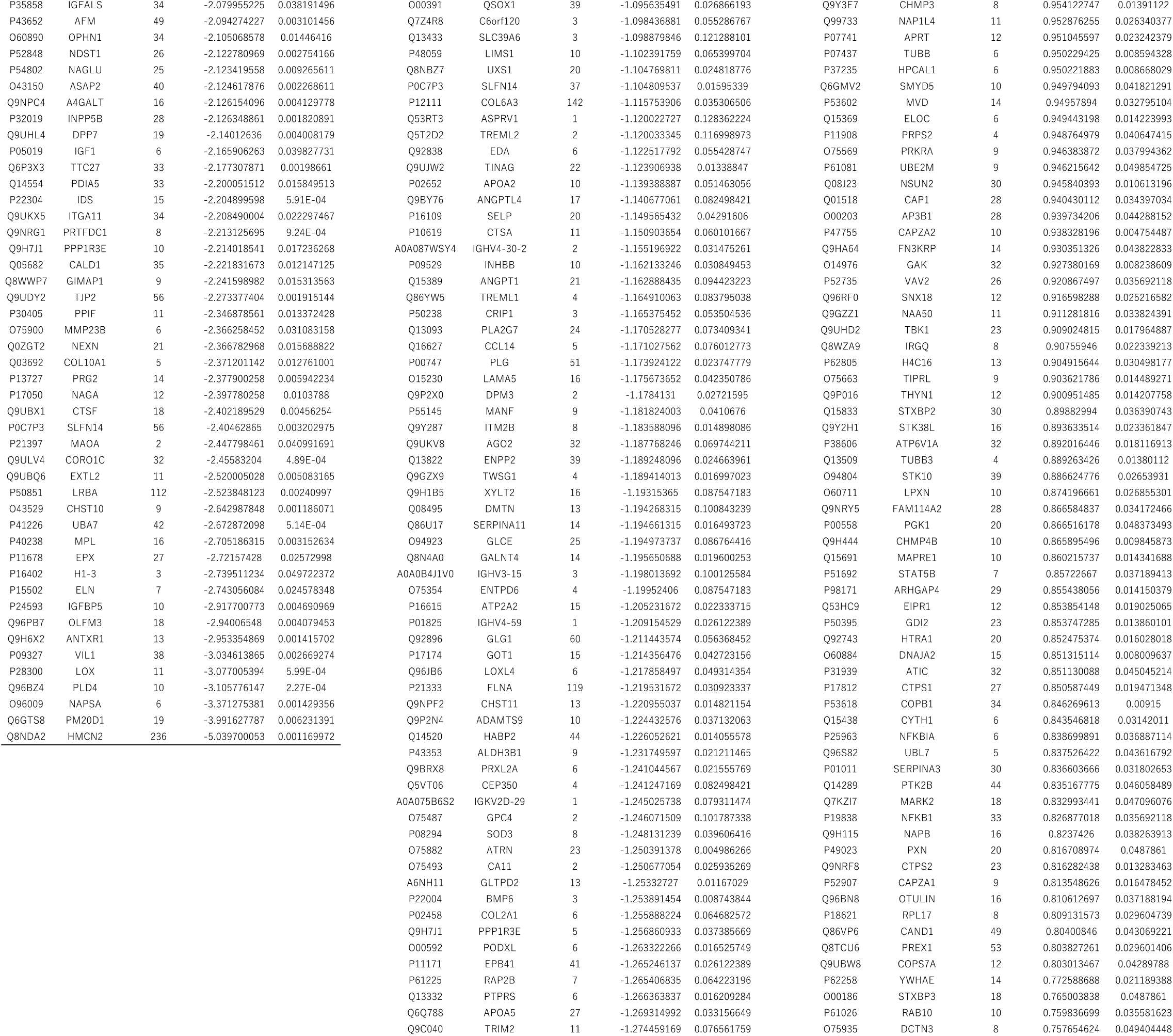
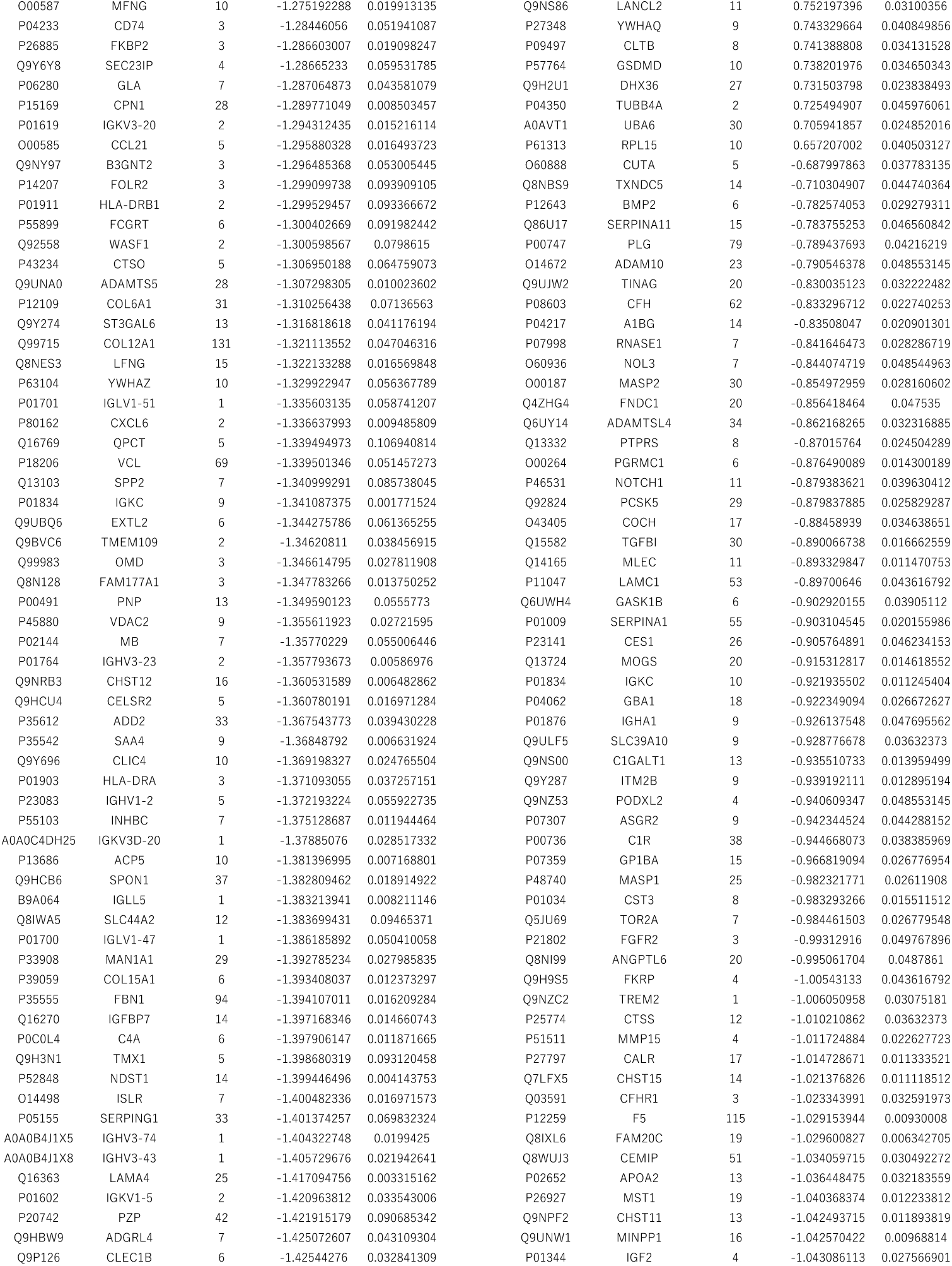
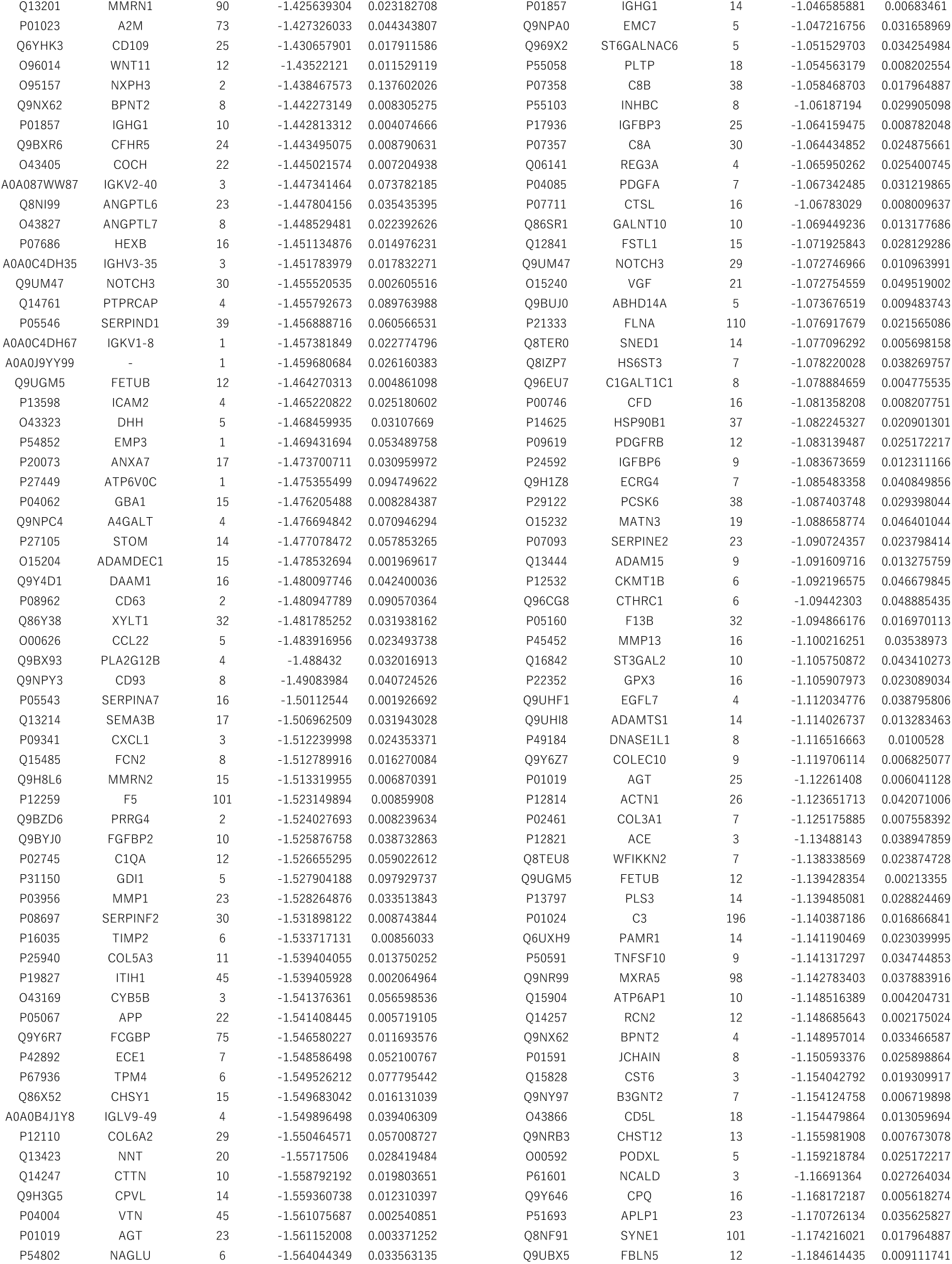
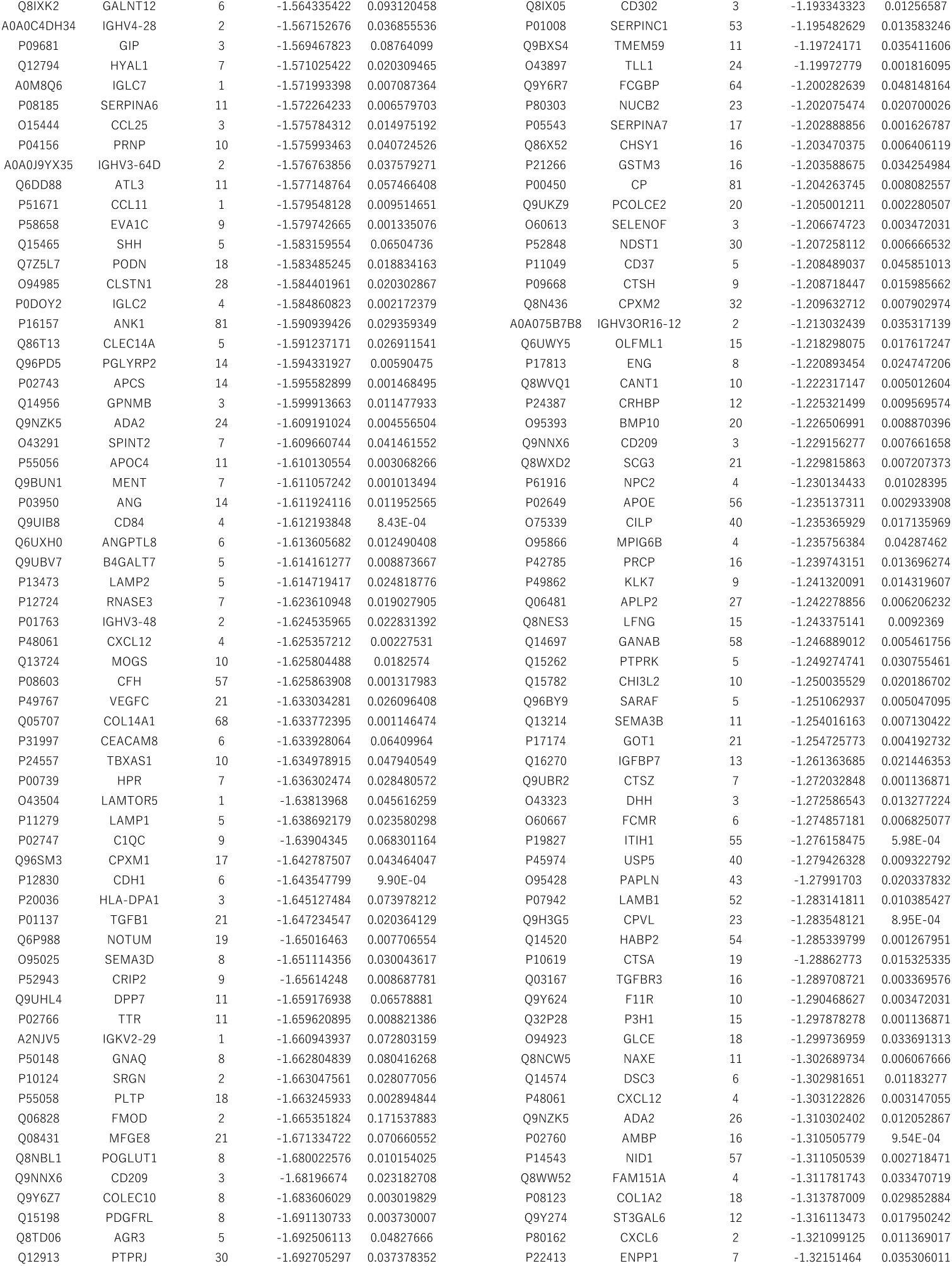
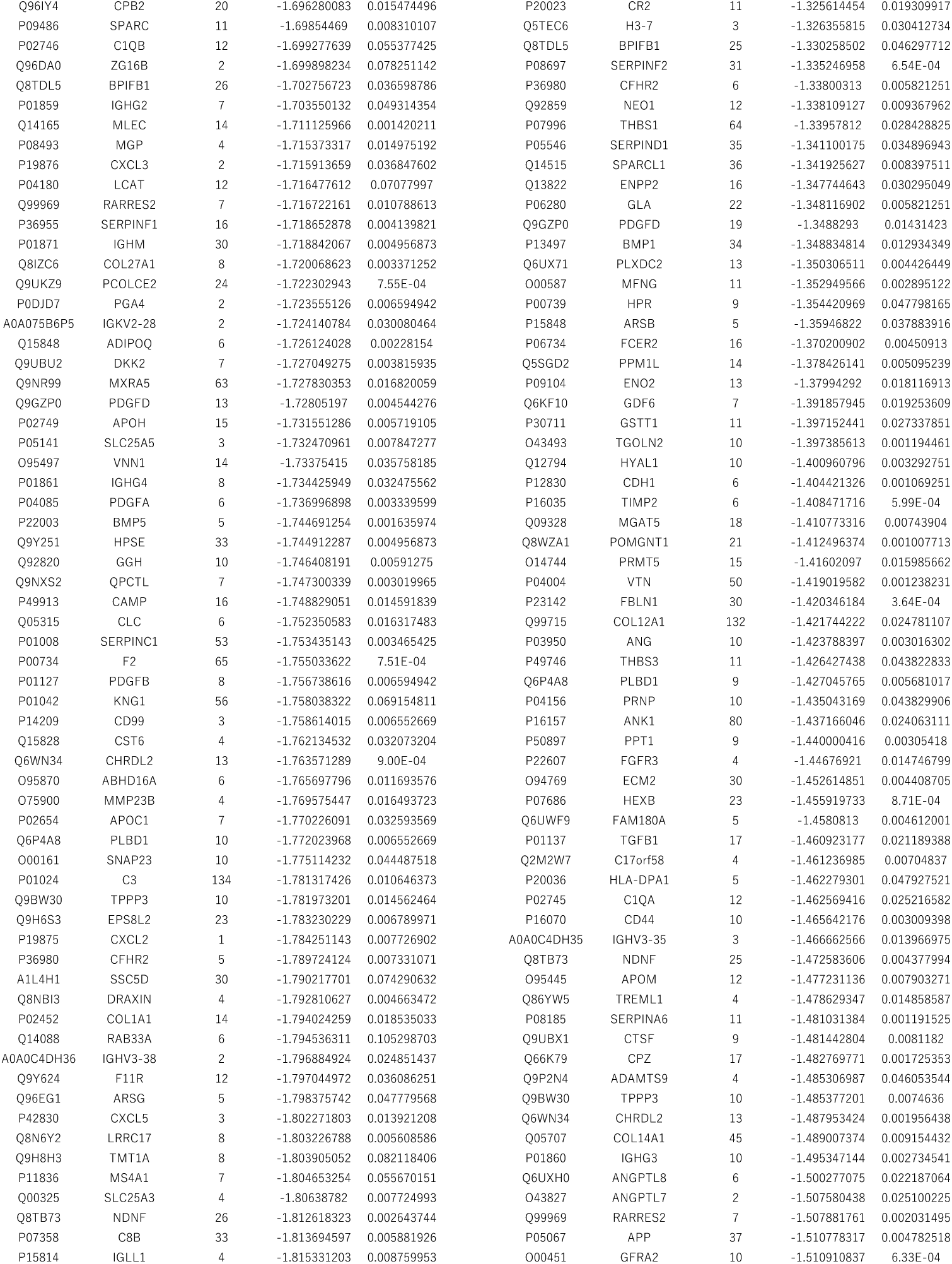
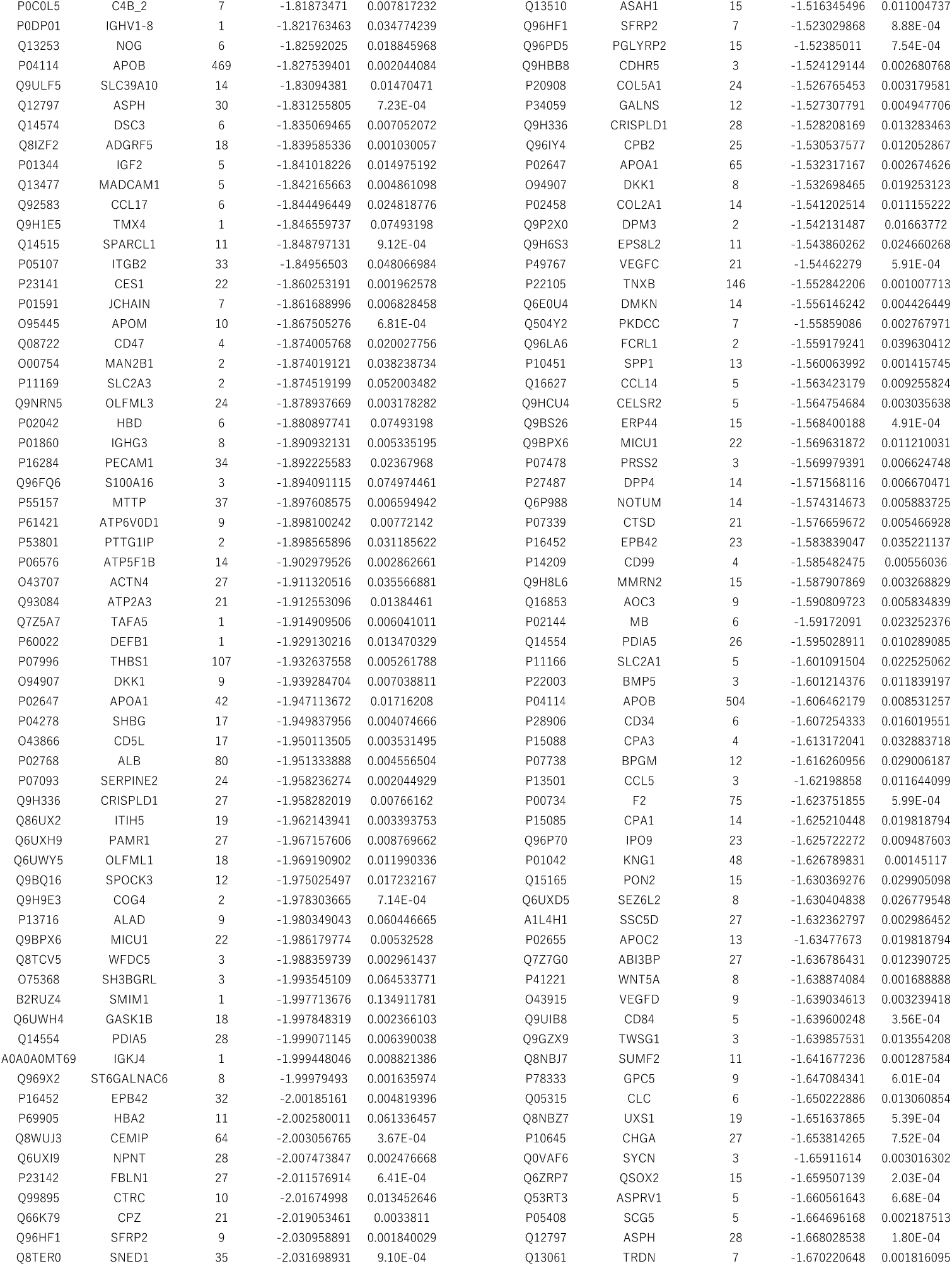
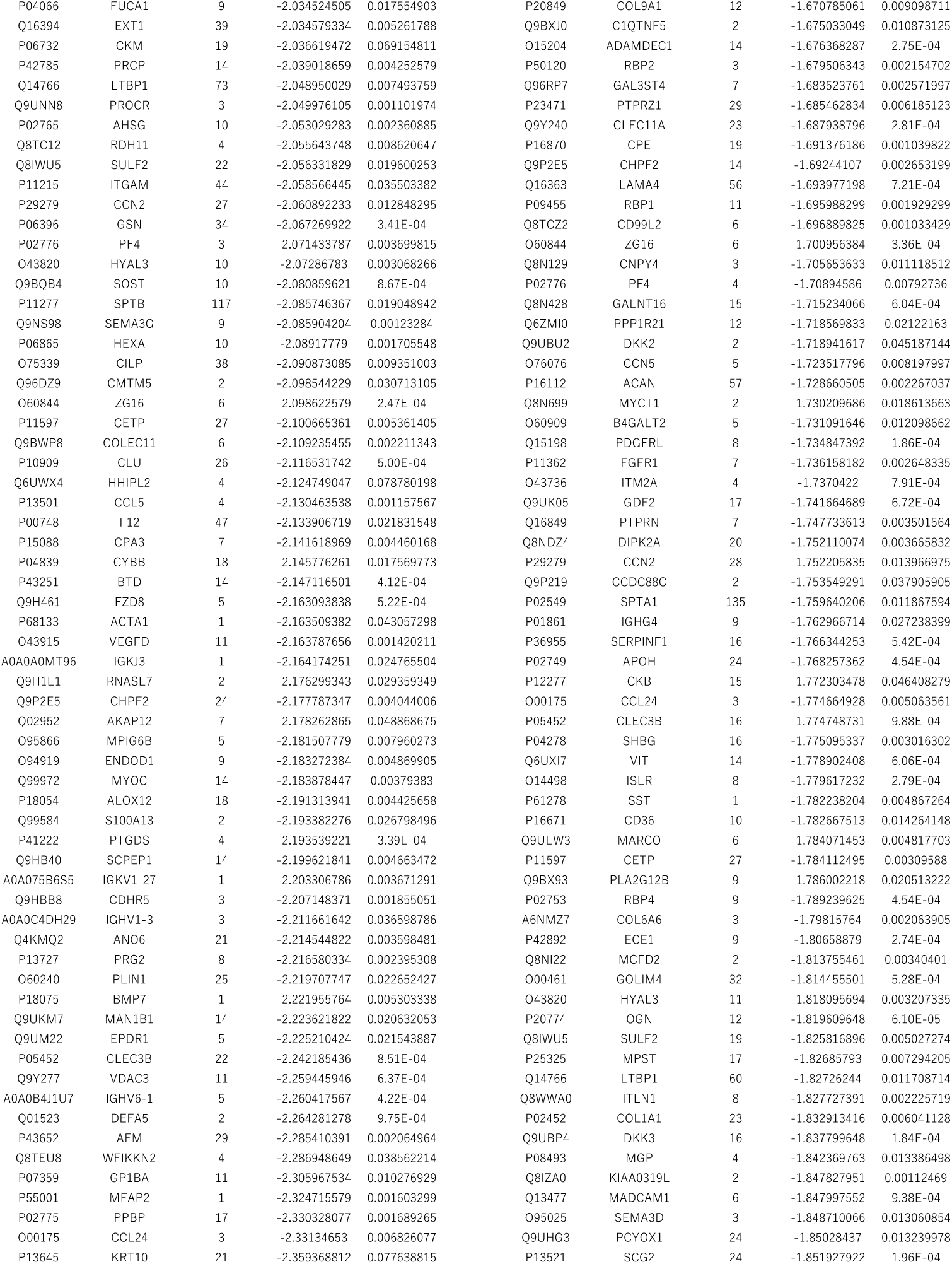
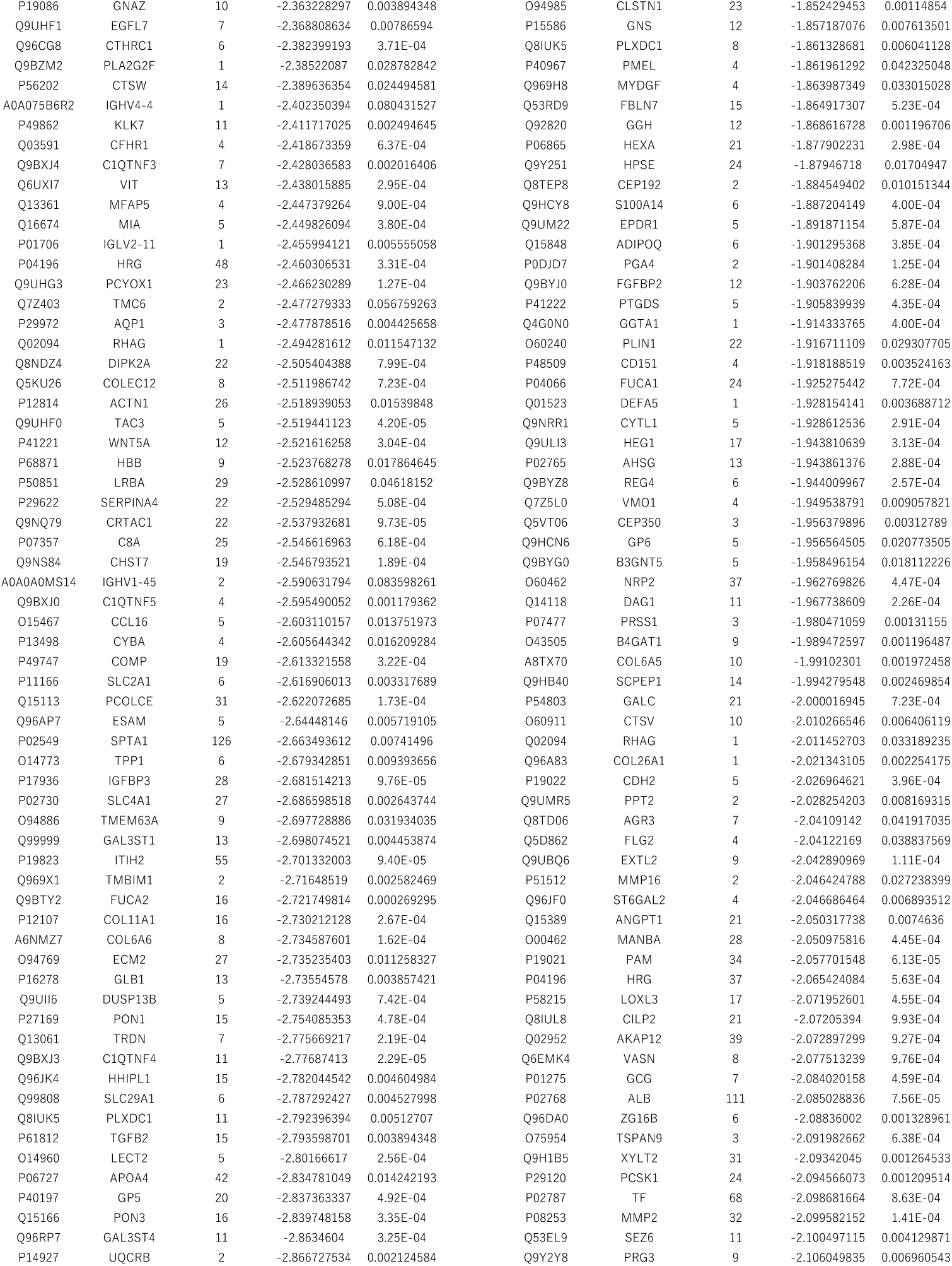
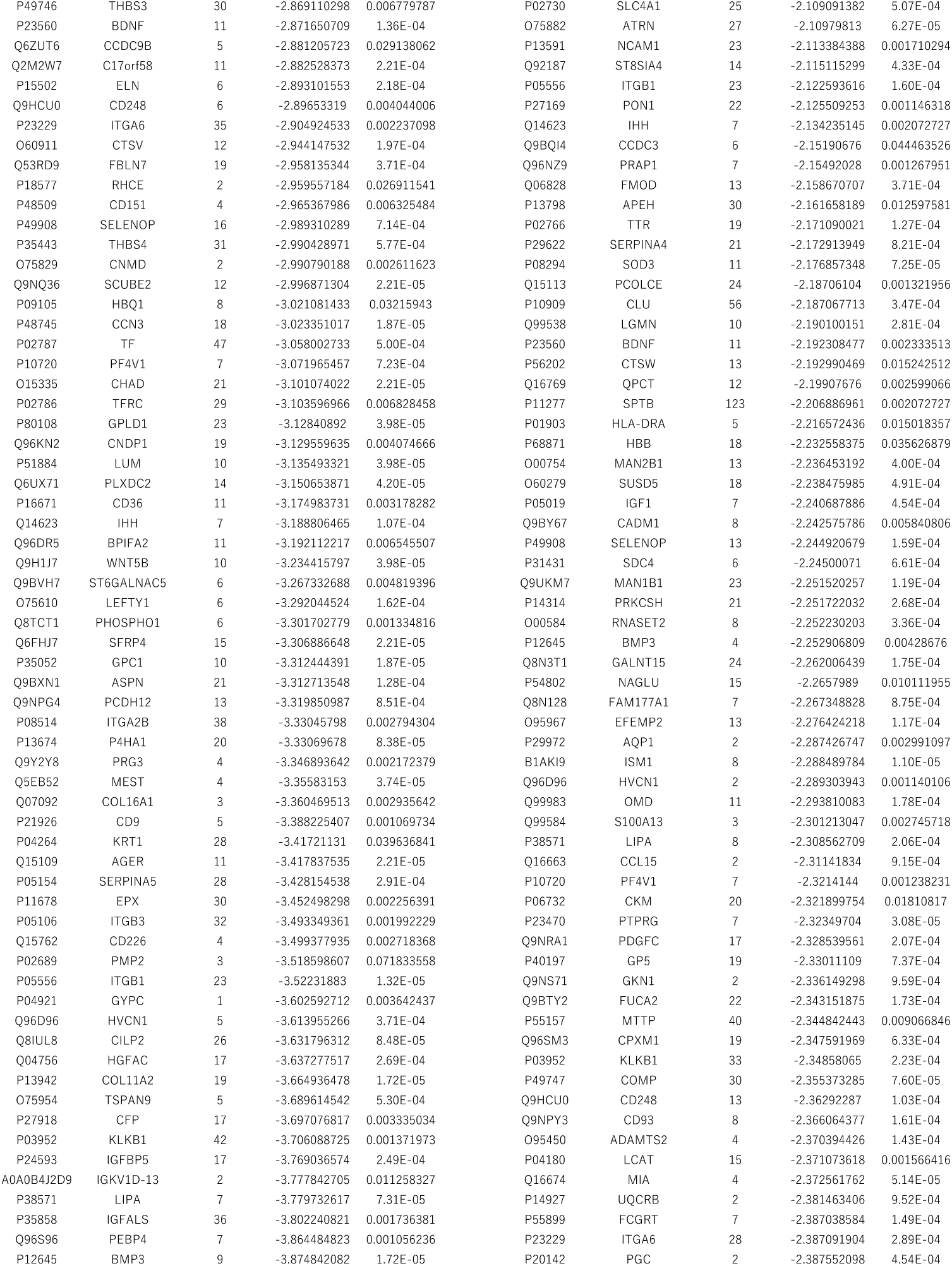
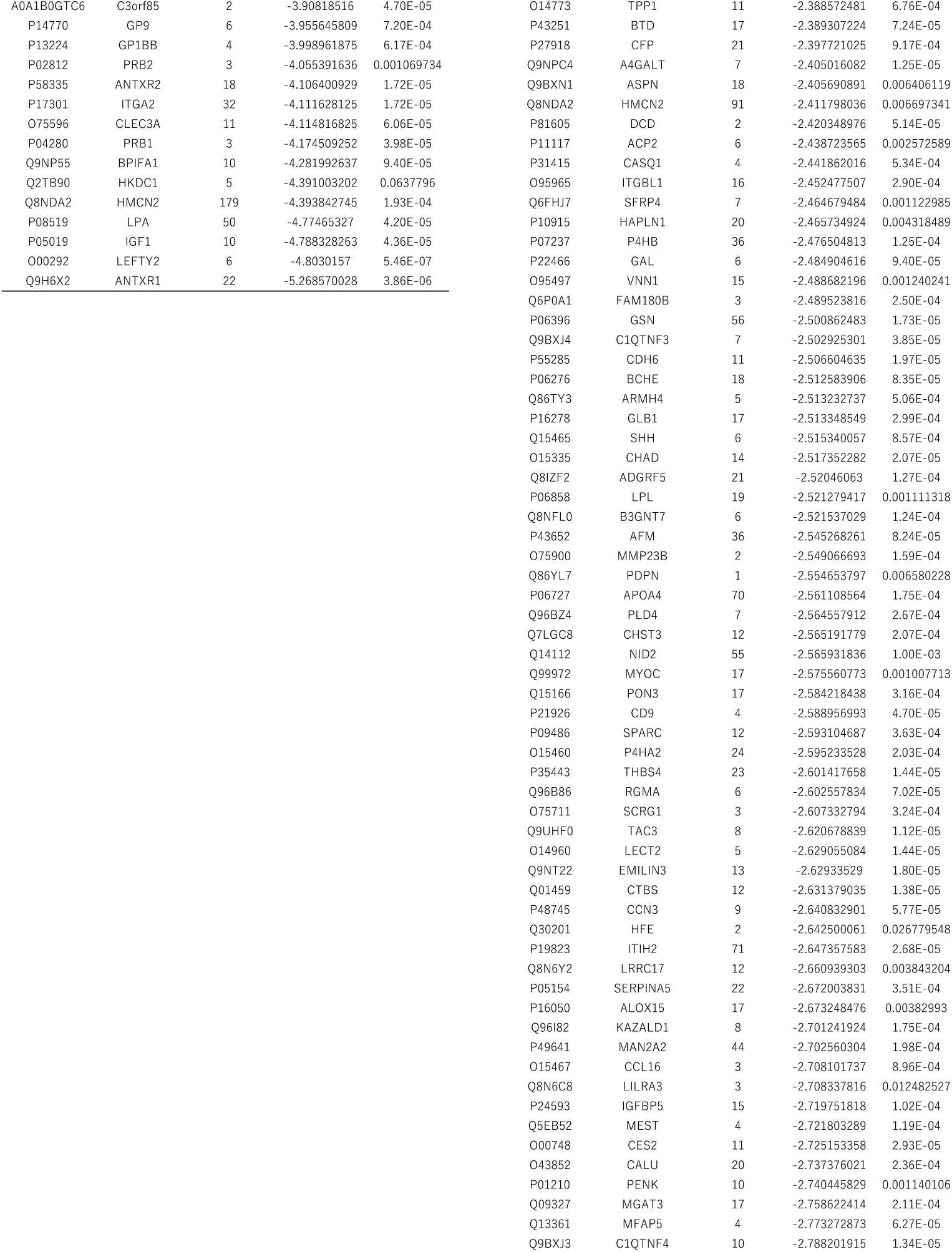
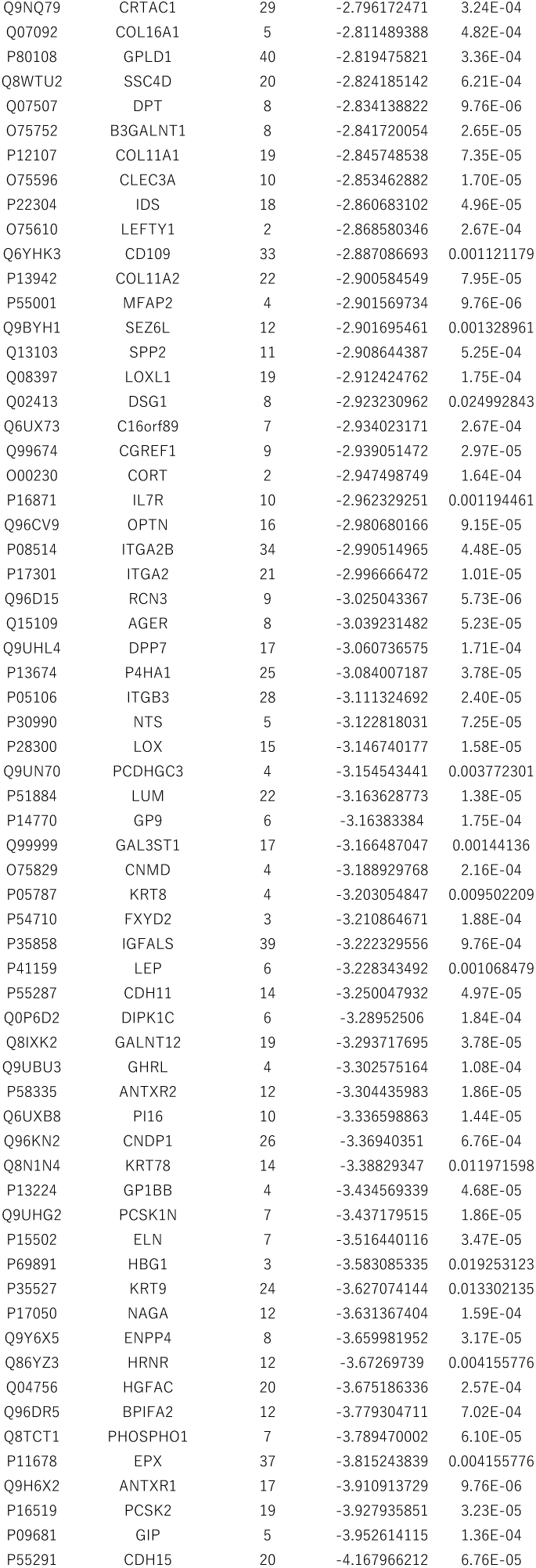
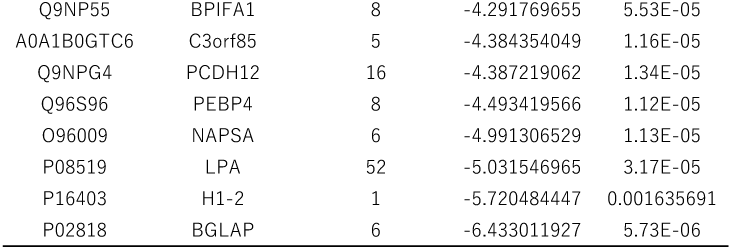
Summary of differentially abundant proteins across four serum preprocessing workflows.

Volcano plots showed that DEPs were detected in all workflows, with both increased and decreased abundance changes observed in patients with sJIA compared with controls (Fig. 2A–D). The distributions of fold change and statistical significance differed across the workflows. Representative inflammation-related biomarkers associated with sJIA, including CRP, ferritin, and IL-18, were upregulated in all the workflows. In contrast, workflow-dependent differences were observed for PTX3, CD163, and serum amyloid proteins, which showed smaller fold changes in some workflows or did not meet the DEP criteria.

**Fig. 2.**
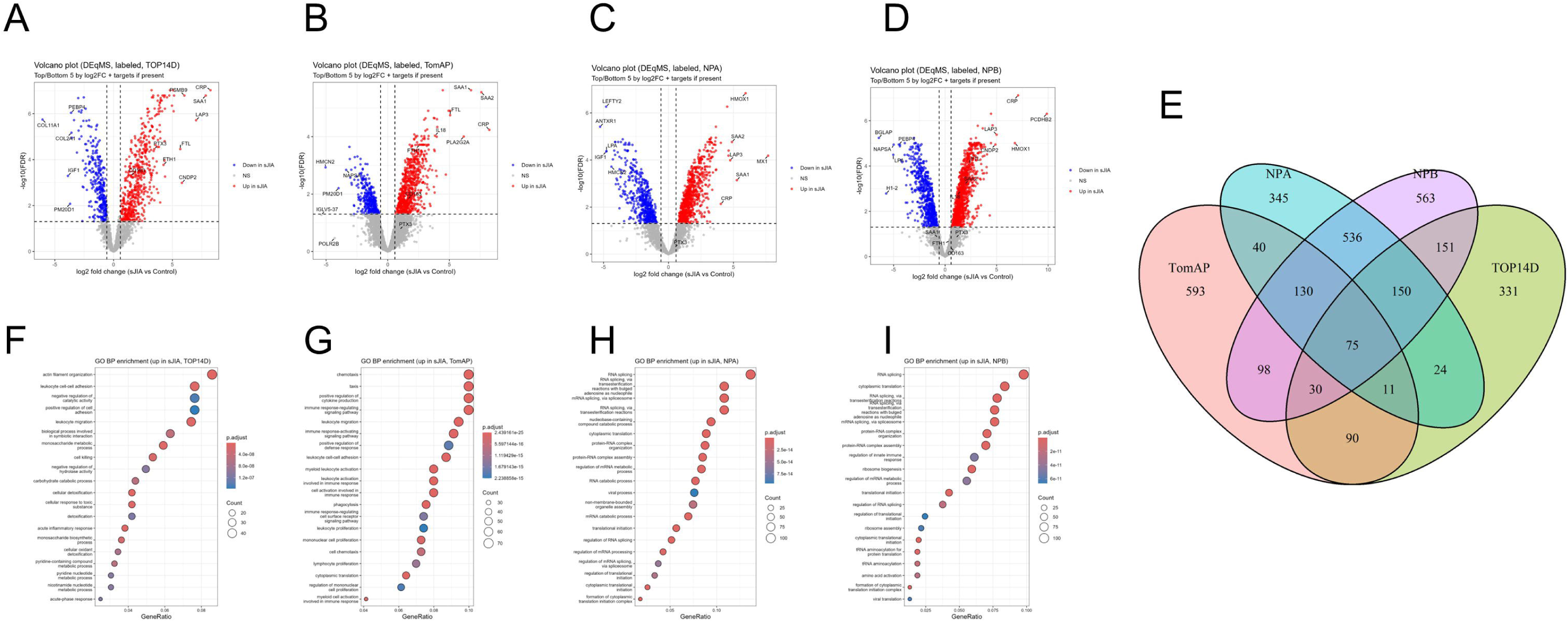
Comparison of sJIA-associated differential protein signatures and functional enrichment patterns across serum preprocessing workflows. A–D. Volcano plots of differential abundance between the sJIA and control groups for TOP14D (A), TomAP (B), NPA (C), and NPB (D). The x-axis represents the log_2_ fold change (log_2_FC) for sJIA versus controls, and the y-axis represents −log10(FDR). Red points indicate DEPs upregulated in sJIA, blue points indicate DEPs downregulated in sJIA, and gray points indicate proteins that did not meet the DEP criteria. Vertical dashed lines indicate log_2_FC = ±0.58, and the horizontal dashed line indicates FDR = 0.05. Proteins with the largest positive and negative log_2_FC values, together with predefined target proteins when detected, are labeled. E. Venn diagram showing the overlap of DEPs identified across the four workflows. Values within each region indicate the numbers of DEPs unique to or shared among the corresponding workflows. Seventy-five DEPs were common to all four workflows. F–I. Over-representation analysis of Gene Ontology Biological Process (GOBP) terms using proteins classified as upregulated in sJIA for TOP14D (F), TomAP (G), NPA (H), and NPB (I). The top-enriched terms are shown. Dot size indicates the number of input genes assigned to each term (Count), and dot color represents the Benjamini–Hochberg-adjusted p-value (p.adjust). DEP, differentially abundant protein; FDR, false discovery rate; GOBP, Gene Ontology Biological Process; log_2_FC, log_2_ fold change; NPA and NPB, nanoparticle-based enrichment workflows; sJIA, systemic juvenile idiopathic arthritis; TomAP, tomato lectin affinity purification; TOP14D, Top14 abundant protein depletion.

Next, we compared the overlap of DEPs obtained from each workflow. DEPs shared across workflows were observed, and workflow-specific DEPs were also detected (Fig. 2E). TomAP and NPB yielded the largest numbers of proteins that met the DEP criteria. A substantial overlap was also observed between NPA and NPB, indicating that the differential abundance signatures were more similar between workflows with related preprocessing designs. Only 75 DEPs were common to all four workflows (Table 3). These results indicate that changes in sJIA-associated protein abundance were detected across workflows, but that many proteins identified as DEPs were workflow-dependent. Although DEP overlap was influenced by the significance threshold, both shared and workflow-specific DEPs were observed.

**Table 3.** Seventy-five deps shared across all four workflows.

| # | Uniprot ID | gene name |
| --- | --- | --- |
| 1 | P02741 | CRP |
| 2 | P13796 | LCP1 |
| 3 | P52790 | HK3 |
| 4 | Q07960 | ARHGAP1 |
| 5 | P80723 | BASP1 |
| 6 | P05109 | S100A8 |
| 7 | P06702 | S100A9 |
| 8 | P08238 | HSP90AB1 |
| 9 | Q9H6X2 | ANTXR1 |
| 10 | Q12906 | ILF3 |
| 11 | Q12905 | ILF2 |
| 12 | P29966 | MARCKS |
| 13 | P51884 | LUM |
| 14 | P18428 | LBP |
| 15 | Q8IUL8 | CILP2 |
| 16 | Q9NUQ9 | CYRIB |
| 17 | P43652 | AFM |
| 18 | Q13177 | PAK2 |
| 19 | Q15113 | PCOLCE |
| 20 | P13639 | EEF2 |
| 21 | P52209 | PGD |
| 22 | P0DMV8 | HSPA1A |
| 23 | P24593 | IGFBP5 |
| 24 | P34931 | HSPA1L |
| 25 | P06396 | GSN |
| 26 | P31949 | S100A11 |
| 27 | P01011 | SERPINA3 |
| 28 | P61160 | ACTR2 |
| 29 | P30050 | RPL12 |
| 30 | O60664 | PLIN3 |
| 31 | Q9BR76 | CORO1B |
| 32 | Q9UJ70 | NAGK |
| 33 | O15144 | ARPC2 |
| 34 | O15511 | ARPC5 |
| 35 | P05154 | SERPINA5 |
| 36 | P61586 | RHOA |
| 37 | P61158 | ACTR3 |
| 38 | P46976 | GYG1 |
| 39 | Q99536 | VAT1 |
| 40 | O15143 | ARPC1B |
| 41 | P02750 | LRG1 |
| 42 | O15335 | CHAD |
| 43 | Q05639 | EEF1A2 |
| 44 | P17301 | ITGA2 |
| 45 | P59998 | ARPC4 |
| 46 | P52848 | NDST1 |
| 47 | Q8NDA2 | HMCN2 |
| 48 | P23381 | WARS1 |
| 49 | Q9BXN1 | ASPN |
| 50 | P24158 | PRTN3 |
| 51 | P54802 | NAGLU |
| 52 | O14960 | LECT2 |
| 53 | P84095 | RHOG |
| 54 | P22626 | HNRNPA2B1 |
| 55 | P35858 | IGFALS |
| 56 | P05019 | IGF1 |
| 57 | P41222 | PTGDS |
| 58 | P05362 | ICAM1 |
| 59 | P80108 | GPLD1 |
| 60 | Q99983 | OMD |
| 61 | P14317 | HCLS1 |
| 62 | Q8IUE6 | H2AC21 |
| 63 | Q6IBS0 | TWF2 |
| 64 | Q96C19 | EFHD2 |
| 65 | Q5JSH3 | WDR44 |
| 66 | Q04756 | HGFAC |
| 67 | O14498 | ISLR |
| 68 | Q01518 | CAP1 |
| 69 | P02766 | TTR |
| 70 | P05062 | ALDOB |
| 71 | O15204 | ADAMDEC1 |
| 72 | P23229 | ITGA6 |
| 73 | P11234 | RALB |
| 74 | O14773 | TPP1 |
| 75 | P23560 | BDNF |

### Functional enrichment of increased proteins differs across serum preprocessing workflows

GO Biological Process (GOBP) and Reactome pathway enrichment analyses were performed on the upregulated proteins identified in the sJIA group for each serum preprocessing workflow. The most enriched biological processes and pathways differed among the workflows. In the GOBP analysis of TOP14D, terms related to cell adhesion, migration, and inflammatory responses, including leukocyte cell-cell adhesion, actin filament organization, acute inflammatory response, and acute-phase response, were among the most enriched (Fig. 2F). In the Reactome analysis, neutrophil degranulation and interleukin signaling were among the top rank pathways, together with pathways related to stress responses and antigen presentation, including the KEAP1–NFE2L2 (NRF2) pathway, ER phagosome, and cross-presentation (Fig. S2A). In the GOBP analysis of TomAP, terms related to immune cell recruitment and activation during inflammation, including chemotaxis, taxis, leukocyte migration, positive regulation of cytokine production, and phagocytosis, were enriched among the top terms (Fig. 2G). In the Reactome analysis, neutrophil degranulation was the top-ranked pathway, and pathways related to translation initiation and nonsense-mediated decay were also identified (Fig. S2B).

In contrast, in NPA and NPB, the top GOBP terms were consistently related to RNA splicing, protein–RNA complex organization, ribosome biogenesis, and mRNA/RNA catabolic processes (Fig. 2H, I). Reactome analysis also highlighted pathways related to RNA processing and translation, including mRNA splicing and the processing of capped intron-containing pre-mRNAs. In NPB, ROBO receptor signaling was identified among the top-ranked pathways (Fig. S2C, D). Overall, the functional enrichment patterns of the upregulated proteins differed across serum preprocessing workflows, with TOP14D and TomAP showing enrichment patterns related mainly to neutrophil/myeloid and inflammatory processes, and NPA and NPB showing enrichment patterns related mainly to RNA processing and translation. These results indicate that even when the same sJIA samples were analyzed, the biological processes and pathways that ranked highly in the enrichment analyses differed depending on the serum preprocessing workflow.

### Coverage and enrichment patterns of curated sJIA-related gene sets differ across serum preprocessing workflows

In addition to standard functional enrichment analyses, we constructed curated sJIA-related gene sets reflecting key biological processes related to the pathophysiology of sJIA and compared their protein coverage and fgsea enrichment patterns across the serum preprocessing workflows (Tables S1–S4). TomAP consistently showed higher coverage of the proteins included in each gene set. In particular, for the Innate TLR/NLR set, TomAP detected 136 of 231 proteins (58.8%), a higher proportion than for TOP14D (21/231, 9.1%), NPA (69/231, 29.9%), and NPB (82/231, 35.5%) (Fig. 3A–D). In the fgsea analysis, the Innate TLR/NLR set showed significant positive enrichment in TomAP (NES = 1.73, FDR = 6.99 × 10^-4). Positive enrichment was also observed for the AcutePhase/Complement, Neutrophil activation, and Macrophage/MAS gene sets in TomAP (NES = 1.72–2.07; FDR = 7.06 × 10^−4^ for all; Fig. 3E and Fig. S3A–D). These results indicate that among the workflows examined, TomAP showed the broadest coverage and positive enrichment of curated sJIA-related gene sets associated with inflammation, innate immunity, and MAS.

**Fig. 3.**
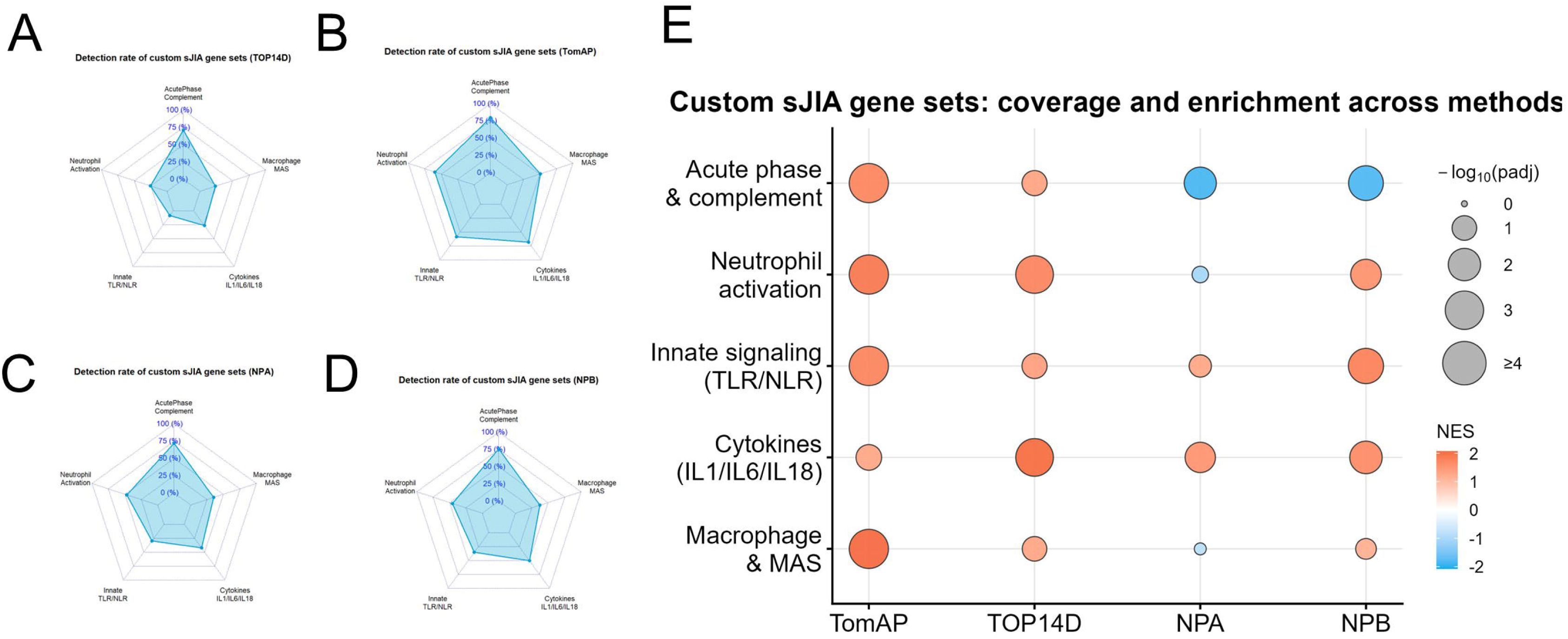
Coverage and enrichment patterns of curated sJIA-related gene sets across serum preprocessing workflows. Differential abundance results from the sJIA (n = 6) and control (n = 6) groups were used to compare the five curated sJIA-related gene sets across workflows. A–D. The protein coverage of each gene set in TOP14D (A), TomAP (B), NPA (C), and NPB (D). Coverage was calculated as the proportion of proteins in each curated gene set detected using the corresponding workflow. E. Pre-ranked GSEA of curated sJIA-related gene sets across workflows. Gene symbol-level log_2_FC values were used as ranking scores for fgsea. Dot color represents the normalized enrichment score (NES), with red and blue indicating positive and negative enrichment, respectively. Dot size represents the −log10-transformed p-value adjusted using the Benjamini–Hochberg method. FDR, false discovery rate; MAS, macrophage activation syndrome; NES, normalized enrichment score; NPA and NPB, nanoparticle-based enrichment workflows; sJIA, systemic juvenile idiopathic arthritis; TomAP, tomato lectin affinity purification; TOP14D, Top14 abundant protein depletion.

In contrast, NPA and NPB showed gene-set-dependent differences in enrichment direction. The Acute phase/complement set showed negative enrichment[in NPA and NPB], whereas the cytokines IL-1/IL-6/IL-18 set showed positive enrichment in TOP14D and NPB (TOP14D: NES = 2.05; NPB: NES = 1.14). Overall, these findings indicate that serum preprocessing workflow selection was associated not only with protein coverage but also with the disease axis readouts captured by curated sJIA-related gene set analysis.

### Differential abundance of sJIA pathogenesis-related proteins across workflows: the inflammasome/interferon axis

Finally, for innate immunity-, inflammasome-, and interferon-related molecules associated with the pathogenesis of sJIA, we compared detectability and abundance changes between the sJIA and control groups across serum preprocessing workflows. Among the proteins in the innate TLR/NLR set, NLRC4, GSDMD, and MEFV were detected and included in the differential abundance analysis of TomAP, NPA, and NPB. However, they were not included in the analysis of TOP14D (Fig. 4A). We further compared the fold changes and DEP classifications for each molecule across the workflows. The detectability and abundance change patterns of inflammasome/interferon-related proteins differed among the workflows. In particular, in TomAP and NPB, sJIA–control differences were evaluated for multiple proteins, including NLRC4, IL-18, PYCARD, OAS3, and MYD88, several of which met the DEP criteria (Fig. 4B). These results indicate that the detection and differential abundance assessment of innate immunity-, inflammasome-, and interferon-related proteins in sJIA are workflow-dependent.

**Fig. 4.**
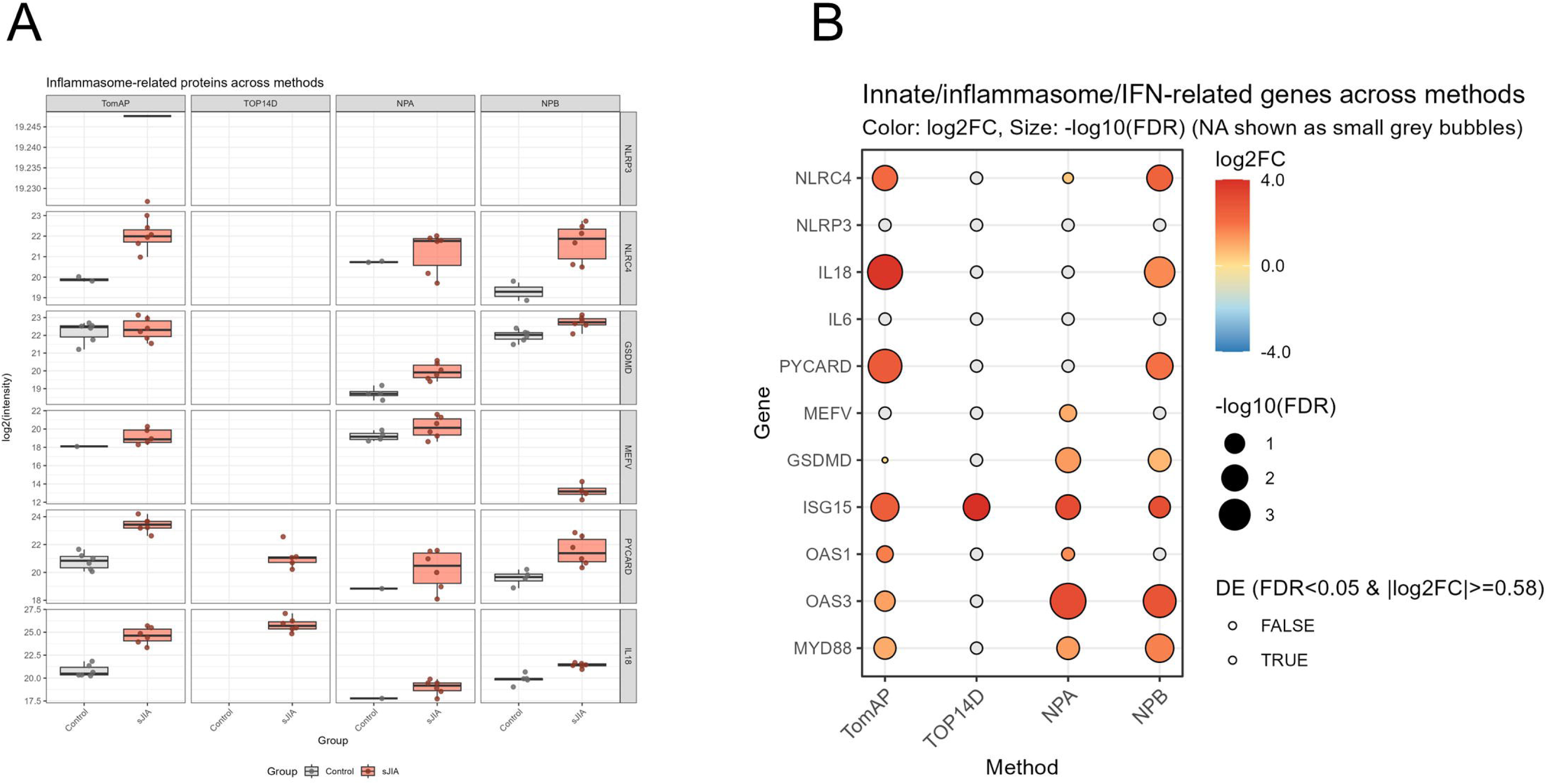
Workflow-dependent detection and differential abundance of innate immune, inflammasome, and interferon-related proteins. Selected innate immune-, inflammasome-, and interferon-related proteins were compared across workflows using samples from the sJIA (n = 6) and control (n = 6) groups. A. Log_2_-transformed protein intensities of NLRP3, NLRC4, GSDMD, MEFV, PYCARD, and IL-18 in the TomAP, TOP14D, NPA, and NPB workflows. Points represent individual samples, boxes represent the interquartile range, and centerlines indicate the median. Only the samples with quantifiable values in the corresponding workflow are shown. B. Bubble plot comparing differential abundance results for selected innate immune-, inflammasome-, and interferon-related proteins across workflows. Color represents the log_2_FC for sJIA versus controls, and bubble size represents −log10(FDR). Proteins not included in the differential abundance analysis are shown as small grey bubbles. The point outlines indicate whether the DEP criteria were met. DEP, differentially abundant protein; FDR, false discovery rate; IFN, interferon; log_2_FC, log_2_ fold change; NPA and NPB, nanoparticle-based enrichment workflows; sJIA, systemic juvenile idiopathic arthritis; TomAP, tomato lectin affinity purification; TOP14D, Top14 abundant protein depletion.

## Discussion

In this study, we systematically compared four serum preprocessing workflows (TomAP, TOP14D, NPA, and NPB) using samples from the same sJIA cohort and a unified DIA-MS acquisition and analysis pipeline. We evaluated how these workflows shaped the detected protein set, differential abundance signatures, and disease axis readouts. Recent studies in plasma proteomics have emphasized that deeper proteome coverage does not necessarily ensure quantitative robustness or biological fidelity, particularly in nanoparticle-based workflows, where preanalytical variation and cellular carry-over require careful consideration (25). Our study extends this depth-versus-fidelity perspective to a disease-oriented setting using sJIA serum and indicates that workflow differences may influence not only proteome depth but also differential abundance signatures and biological readouts. These findings support the view that serum preprocessing is not merely a preparatory step but an important component of experimental design that can shape which disease axes are preferentially captured.

The main findings of this study can be summarized as follows. First, the proteome depth and composition of the detected protein set differed substantially across the workflows. Second, although the sJIA and control groups were separated in all workflows, the overlap of DEPs was limited, and the differential abundance signatures were strongly workflow-dependent. Third, these differences extended to functional interpretations: TOP14D and TomAP preferentially captured enrichment patterns related to inflammatory, neutrophil/myeloid, and cytokine-associated processes, whereas NPA and NPB captured RNA processing- and translation-related signals more prominently. Thus, even when the same disease and serum samples were analyzed, the major biological readouts differed according to the preprocessing workflow.

This distinguishes the present study from several previous workflow comparison studies. Recent comparisons of depletion and enrichment workflows in plasma and serum proteomics have clarified technical characteristics, such as proteome depth, repeatability, and quantification bias (12). However, many of these studies were based on healthy donors or technical replicates, and the extent to which workflow differences influence disease-control comparisons and pathway-level interpretations in a specific disease context remains poorly explored. By applying multiple workflows to sJIA, a systemic autoinflammatory disease, our study adds a disease-oriented benchmark to existing technical comparisons and shows how workflow-dependent differences can propagate into biological interpretation.

Another important finding was the limited overlap of DEPs across the workflows. Only a small subset of DEPs was shared by all four workflows, consistent with the presence of a core sJIA-associated signature detected across the preprocessing strategies. Simultaneously, several differentially abundant signals were detected in a workflow-dependent manner. This likely reflects differences in the detected protein set and the quantitative characteristics of each workflow, which can yield distinct differential abundance signatures even when the same disease samples are analyzed. Therefore, differences among workflows should not be interpreted solely as differences in the number of detected proteins but also as differences in which aspects of disease biology are more readily captured. In serum proteomics, preprocessing should be considered an explicit source of analytical variation.

This workflow dependency was particularly evident in functional analyses. In TOP14D and TomAP, terms and pathways related to neutrophil/myeloid inflammation, cell migration, and cytokine responses were among the most enriched. TomAP showed broad coverage and positive enrichment for curated sJIA-related gene sets, including Innate TLR/NLR, Neutrophil activation, and macrophage/MAS, suggesting that this workflow may be useful for identifying inflammation-, innate immunity-, and MAS-related biology in sJIA serum. In contrast, NPA and NPB highlighted modules related to RNA splicing and translation, suggesting that these nanoparticle-based workflows may be useful for the exploratory profiling of protein classes that are less prominent in other workflows. However, these findings should be interpreted with caution. RNA processing- and translation-related signals may reflect aspects of sJIA biology, but they may also reflect protein classes preferentially recovered by these workflows or the influence of preanalytical cellular carry-over. Recent plasma proteomic studies have raised the depth-versus-fidelity issue in nanoparticle-based workflows (25). Because this study used serum, plasma-based contamination frameworks cannot be directly applied; the NPA/NPB findings should therefore be interpreted with both biological signals and workflow-dependent biases in mind.

The practical implication of this study is that serum preprocessing workflows should be selected according to the disease axis or analytical objective, rather than solely based on proteome depth. By combining general GO/Reactome analyses with curated sJIA-related gene sets, we compared workflow-dependent biological readouts from a disease axis perspective. In this context, TomAP may be suitable for assessing inflammation, innate immunity, and MAS-related biology, whereas NPA and NPB may be useful for broader exploratory proteomic profiling. TOP14D and NPB may also provide complementary information on cytokine-related axes. Importantly, our results do not identify a single universally superior workflow; rather, they support the need for a fit-for-purpose workflow selection according to the research objective.

Workflow dependence has also been observed in the detection of inflammasome/interferon-related proteins. Molecules such as IL-18, PYCARD, NLRC4, GSDMD, and MEFV showed different detection patterns and differential abundance behaviors across workflows, indicating that the apparent readout of the inflammasome/IL-18 axis may depend on serum preprocessing. Because innate immune cytokines and MAS-related inflammatory pathways are central to the pathophysiology of sJIA, cross-workflow comparisons of these molecules provide information beyond a simple depth comparison and may inform biomarker-oriented study design and the evaluation of disease-related hypotheses.

This study had some limitations. First, the study was exploratory in scale and did not allow conclusions to be drawn regarding the universal superiority or inferiority of any workflow. Second, the cohort was cross-sectional and included patients receiving treatment. Therefore, the observed signals may have been influenced by both the disease activity and treatment status. Third, the control group consisted of only healthy controls, and differentiation from other inflammatory diseases was not evaluated. In addition, this study compared end-to-end workflows that differed in terms of input, chemistry, and post-enrichment processing, and was not a head-to-head comparison in which a single factor was strictly controlled. Future studies using larger independent cohorts, untreated patients, patients with MAS, and inflammatory disease controls and parallel serum/plasma comparisons are required to validate these findings.

## Conclusion

This study indicates that differences in serum preprocessing workflows can shape the disease axis readouts captured by DIA-MS in sJIA serum. These findings support the view that workflow selection in serum proteomics is not merely a technical decision but an important component of the study design that influences which aspects of disease biology are preferentially detected. This disease-oriented benchmark provides practical guidance for fit-for-purpose workflow selection and the design and interpretation of serum proteomics studies, particularly in autoinflammatory diseases.

## Abbreviations

sJIA: systemic juvenile idiopathic arthritis;
MAS: macrophage activation syndrome;
DIA: data-independent acquisition;
DDA: data-dependent acquisition;
TomAP: Tomato lectin affinity purification;
TOP14D: Top14 depletion;
NPA/NPB: nanoparticle-based enrichment fraction A/B;
DEP: differentially expressed protein;
GSEA: gene set enrichment analysis;
GO: Gene Ontology;
GOBP: Gene Ontology Biological Process;
FDR: false discovery rate;
IFN: interferon.

## Acknowledgements

We would like to thank Editage (www.editage.jp) for English language editing and their assistance in creating the graphical abstract for this paper.

## Author contributions: CRediT

**Hironori Sato**: Conceptualization, Formal analysis, Writing - Original Draft. **Shinji Akioka:** Resources. **Yusei Okuda**: Investigation. **Ryo Konno**: Formal analysis, Investigation. **Yusuke Kawashima**: Investigation, Formal analysis. **Osamu Ohara**: All authors read, revised, and approved the final version of the manuscript.

## Declaration of generative AI use

During the preparation of this work, the authors used ChatGPT (OpenAI; GPT-5.1, accessed November 2025) to assist in the development and debugging of the R code for the DEqMS-based analyses, preliminary figure planning, and English language editing. All the AI-assisted outputs were reviewed, revised, and verified by the authors. All analyses were performed using the underlying data, and the final figures were generated from the original data using reproducible author-verified R scripts. The authors take full responsibility for the content of the published article.

