## Supplemental Figures and Tables for "Serum preprocessing workflows differentially shape biological readout in data-independent acquisition proteomics of systemic juvenile idiopathic arthritis": Supplementary_material_preprint.pdf

### Supplemenatry Material

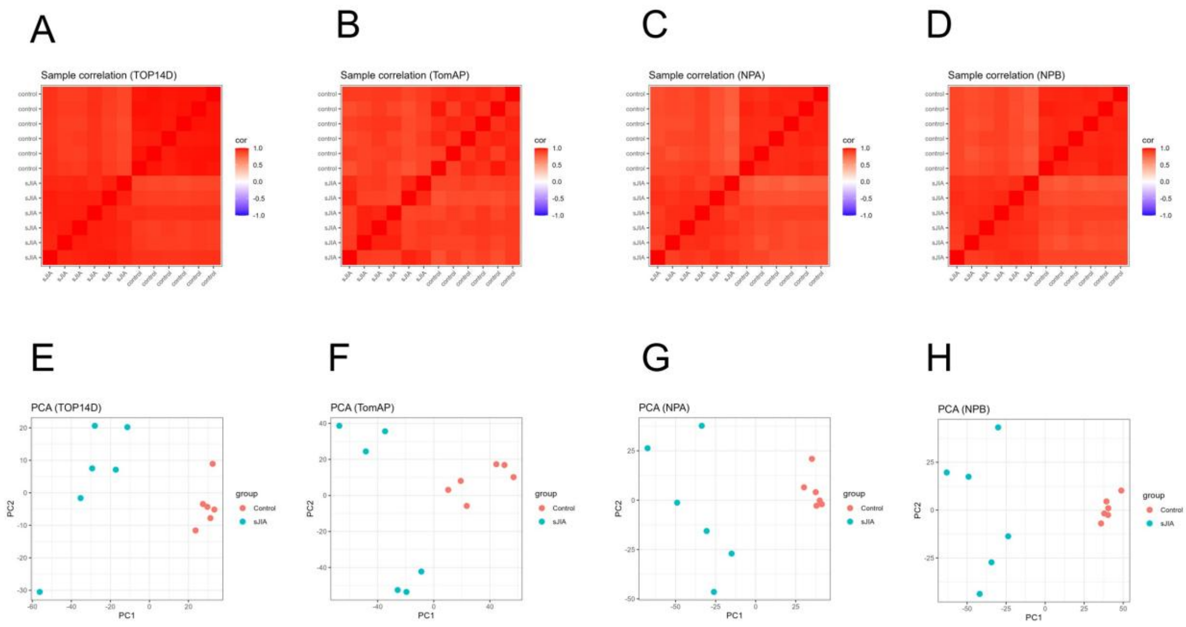


Fig. S1. **Sample Correlations and Principal Component Analysis across Serum Preprocessing Workflows**

Protein intensity data from sJIA (n = 6) and control (n = 6) groups were compared separately for each workflow.

A–D. Inter-sample correlation heat maps for TOP14D (A), TomAP (B), NPA (C), and NPB (D). The color represents the correlation coefficient between each pair of samples.

E–H. Principal component analysis (PCA) for TOP14D (E), TomAP (F), NPA (G), and NPB (H). Each point represents an individual sample, and the colors indicate the sJIA and control groups.

NPA and NPB, nanoparticle-based enrichment workflows; PCA, principal component analysis; sJIA, systemic juvenile idiopathic arthritis; TomAP, tomato lectin affinity purification; TOP14D, Top14 abundant protein depletion.

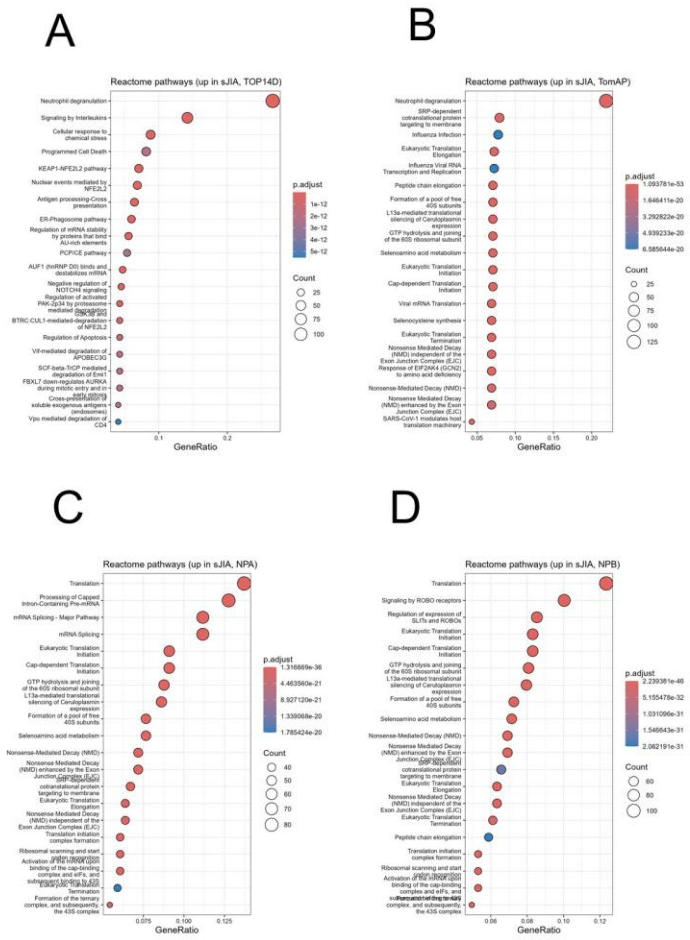


Fig. S2. **Reactome Pathway Enrichment across Serum Preprocessing Workflows**

Reactome pathway over-representation analysis using proteins classified as upregulated in sJIA is shown for TOP14D (A), TomAP (B), NPA (C), and NPB (D). The x-axis represents the GeneRatio, the dot size indicates the number of input genes assigned to each pathway (Count), and the dot color represents the Benjamini–Hochberg-adjusted p-value (p.adjust).

NPA and NPB, nanoparticle-based enrichment workflows; sJIA, systemic juvenile idiopathic arthritis; TomAP, tomato lectin affinity purification; TOP14D, Top14 abundant protein depletion.

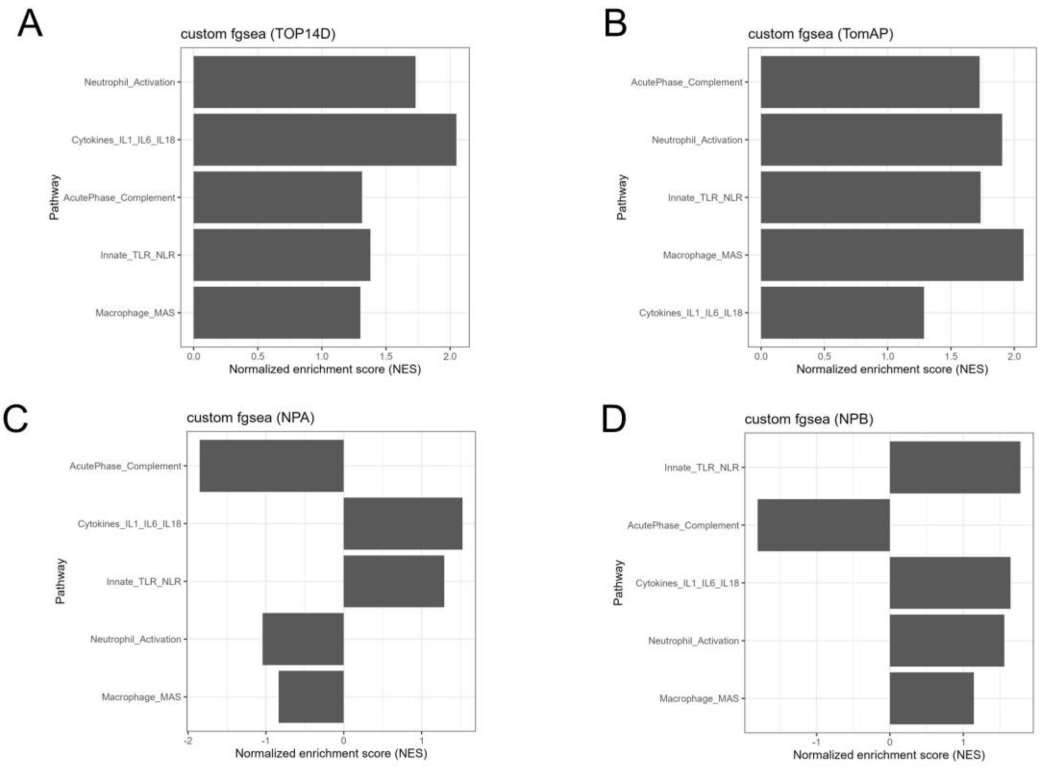


Fig. S3. **Pre-ranked GSEA of Curated sJIA-Related Gene Sets across Serum Preprocessing Workflows**

Pre-ranked GSEA results for the curated sJIA-related gene sets for TOP14D (A), TomAP (B), NPA (C), and NPB (D). The x-axis represents the normalized enrichment score (NES), with positive and negative values indicating enrichment toward higher and lower protein abundances, respectively, in sJIA.

GSEA, gene set enrichment analysis; MAS, macrophage activation syndrome; NES, normalized enrichment score; NPA and NPB, nanoparticle-based enrichment workflows; sJIA, systemic juvenile idiopathic arthritis; TomAP, tomato lectin affinity purification; TOP14D, Top14 abundant protein depletion.
